# Ancient mitogenomes reveal unexpected diversity in Madagascar’s extinct hippopotamuses

**DOI:** 10.64898/2026.10.02.756093

**Authors:** A.T. Salis, E. Hekkala, L.R. Godfrey, D.A. Burney, Z.T. Calamari, A. Caragiulo, N.J. Rahantaharivao, L. Ranivoharimanana, H.A.M. Rasolonjatovo, B.T. Smith, J. Gatesy, C.J. Raxworthy, J.J. Flynn

## Abstract

Madagascar’s extinct hippopotamuses were a distinctive component of the island’s recently lost megafauna, yet their origins, diversification history, and taxonomy have remained unresolved. Here, we recover the first ancient mitochondrial genomes from Malagasy hippopotamuses, obtaining phylogenetic data from 37 subfossil specimens across seven localities and all three named species, and integrate these data with morphology and total-evidence phylogenetic analyses. The 35 endemic specimens form a single clade most closely related to *Hippopotamus amphibius*, supporting a single colonization of Madagascar. Within this radiation, five deeply divergent and geographically structured mitochondrial lineages diversified across most regions of Madagascar during the Pleistocene. Tip-dating estimates a remarkably young crown age of ∼1.41 Ma, following divergence from mainland *Hippopotamus* at ∼3.24 Ma. These lineages do not map cleanly onto traditional taxonomy: material assigned to *H. lemerlei* spans several divergent clades, while *H. laloumena* and *H. madagascariensis* are nested within this broader diversity. Morphology supports some lineage assignments but also points to previously unrecognized diversity, while ancestral-state reconstruction supports early body-size reduction followed by a secondary increase in *H. laloumena*. Malagasy hippopotamuses therefore represent an exceptionally recent radiation of large-bodied island mammals (and the youngest in Madagascar), showing that Madagascar continued to generate endemic mammalian diversity well into the Pleistocene. Their late-Holocene extinction erased the product of more than a million years of rapid insular diversification, the drivers of which remain enigmatic.

## INTRODUCTION

Recent extinctions can erase not only known species, but also cryptic diversity, unrecognized evolutionary lineages, population structure, and ecological breadth that is difficult to recognize from morphology alone. Madagascar provides an exceptional example, having experienced one of the youngest and most dramatic losses of endemic Quaternary fauna. All endemic terrestrial vertebrate species >10 kg went extinct within the last millennium, including up to four elephant bird species, two giant tortoises, three hippopotamus species, two large carnivoran (giant fosa) species, two aardvark-like species, the horned crocodile, and 17 species of lemur^1–4^. Notably, nearly all extinct early-Holocene megafaunal taxa were still present on Madagascar at the time of human arrival several thousand years ago. Although the relative roles of anthropogenic and environmental change remain debated, Madagascar’s megafaunal extinctions are generally understood to reflect a combination of human impacts and natural environmental change^1,3–5^.

The long isolation of Madagascar from other major landmasses, its high levels of endemism, and recent timing of Quaternary megafaunal extinctions present an unusually powerful system for using ancient DNA (aDNA) to investigate the evolution, ecology, systematics, and life history of lost megafauna^5^. To date, only limited ancient DNA research has touched upon the taxonomy, systematics, and biogeography of extinct Malagasy megafauna including the horned crocodile^6^, elephant birds^7^, lemurs^8,9^, and ancient tortoises^10^. aDNA—DNA extracted from archaeological or palaeontological specimens—provides a direct means of recovering evolutionary relationships and genetic structure in extinct taxa, including diversity that may be difficult to diagnose from morphology alone. Most aDNA studies remain biased toward permafrost due to a close relationship between increased ambient temperature and moisture and DNA degradation, making long-term DNA preservation challenging in warmer tropical regions^8,11^. However, advances in high-throughput sequencing and library preparation are increasingly extending aDNA research further back in time and into warmer, more tropical regions^8,11–13^. Madagascar therefore represents both an important and a technically challenging setting in which to apply aDNA to its recently extinct megafauna. The extinct Malagasy hippopotamuses^14–18^ provide an opportunity to use aDNA to test whether current morphology-based systematics captures the evolutionary diversity lost during Madagascar’s late Quaternary extinctions, as well as estimate divergence dates, and confirm new locality records.

The origin of Madagascar’s endemic mammal diversity is understood to have been shaped by dispersal following the island’s isolation rather than by vicariance^19^. Phylogenetic work supports African origins for many Malagasy vertebrate groups, often involving few colonization events followed by in situ diversification^19–21^. For example, each of the four extant Malagasy mammalian groups—lemurs, euplerid carnivorans, tenrecs, and nesomyine rodents—are monophyletic in comparison to African counterparts, likely the result of single dispersal events^22–25^. Rafting is largely considered the main dispersal mechanism but remains contentious with the possibility of dispersal via land bridges^26,27^. In contrast, the mode, timing and number of colonization events underlying the evolutionary history of extinct Malagasy dwarf hippopotamuses remains _unresolved_15,26,28,29.

All Malagasy hippos are recently extinct. Three species of *Hippopotamus* have been described from Pleistocene and Holocene subfossil deposits: 1) *H. lemerlei*, generally associated with rivers, wetlands, and estuaries of western and southern Madagascar^15,18,30^; 2) *H. madagascariensis,* principally in the central highlands and usually interpreted as more terrestrial and robust^15,17^; and 3) *H. laloumena*, a larger form known from eastern coastal Madagascar^16^. Their distinction is based primarily on cranial, dental and postcranial morphology, although incomplete specimens and overlapping characters have complicated species assignment^14,15,31,32^.

The evolutionary origins of Malagasy hippopotamuses have remained uncertain since their discovery. Morphological differences among the different species on the island have been interpreted as evidence of either in situ diversification within Madagascar or as the result of multiple colonisations from Africa^26,33^. Under the latter scenario, different Malagasy species have sometimes been linked to different mainland lineages. Although originally placed within *Hippopotamus*^17^*, Hip. madagascariensis* has been referred to as *Hexaprotodon*, with affinity to the extant West African pygmy hippopotamus, *Choeropsis liberiensis* (previously *Hexaprotodon liberiensis*)^33,34^. The largest of Madagascar’s extinct endemic hippos, *Hip. laloumena* has been proposed to represent an independent colonization by a lineage close to, or potentially indistinguishable from, *Hip. amphibius*^16,35,36^. Consequently, some biogeographic syntheses have invoked up to three separate hippopotamid colonisations of Madagascar^33,36^. No successful DNA retrieval from Malagasy hippopotamus subfossils has been published previously, leaving the relationships, colonization history, and evolutionary diversity of the group to be inferred exclusively from morphology.

Morphological and isotopic data provide essential context for interpreting genetic structure in Malagasy hippopotamuses. Morphological studies have proposed cranial, mandibular, dental, and postcranial characters to distinguish named species, but many specimens are incomplete, sample sizes are small, measurement ranges overlap, and fragmentary material can be difficult to assign^15,31,32^. Stable isotope studies suggest that Malagasy hippopotamuses used a range of habitats and diets, with isotopic variation often differing substantially among regions^37–41^. Integrating genetic, morphological, and isotopic evidence therefore may help determine whether different mitochondrial (mt) lineages correspond to named taxa, geography, diagnostic morphology, or ecological differentiation.

Here, we generate the first mt genome data from subfossil Malagasy hippopotamuses sampled across seven localities and 58 samples representing all three described species. We use this dataset to: 1) test whether Malagasy hippopotamuses represent a single endemic mtDNA radiation or multiple colonization/dwarfing events by including appropriate outgroups; 2) estimate when they diverged from mainland hippopotamids and diversified within Madagascar; and 3) assess whether mtDNA lineages correspond to named species, geography, morphology, or ecological variation. By integrating ancient mitogenomes with morphological measurements and stable isotope data, we then evaluate how much evolutionary and ecological diversity was lost during the recent extinction of Malagasy hippopotamuses.

## RESULTS

### Mitochondrial (mt) DNA recovery from Malagasy hippopotamus subfossils

Of 58 Malagasy hippopotamid specimens processed, 40 (69%) yielded mitochondrial DNA reads uniquely identifiable as hippopotamid. Of these, 37 specimens met our minimum coverage criteria for phylogenetic analysis, with ≥50% mitogenome recovered at ≥3× depth. A reduced high-coverage dataset that included 26 specimens with ≥75% of the mitogenome recovered at ≥3× depth was used for haplotype-network and Bayesian tip-dating analyses.

### Phylogenetic structure of Malagasy hippopotamuses

Maximum-likelihood analyses recovered the same major topology under both outgroup treatments, using *Choeropsis liberiensis* (Figure 1) or the more distant blue whale, *Balaenoptera musculus* (Figure S2). The *B. musculus*-rooted analysis placed the Malagasy radiation closer to *H. amphibius* and the extinct Cypriot insular dwarf, *H. minor*, than to *C. liberiensis* (Figure S2). Hereafter, results primarily refer to the tree rooted with *C. liberiensis*. Of the 37 Malagasy hippopotamid samples with sufficient DNA, 35 formed a monophyletic endemic clade (Figure 1). The remaining two specimens, both originally identified as *H. laloumena*^1^, grouped with modern African mainland *H. amphibius*. Their lack of provenance, very young radiocarbon ages and morphology are consistent with the interpretation by Faure, et al. ^14^ that they were historical specimens brought from mainland Africa to the Academy Malgache (Antananarivo, Madagascar) to serve as comparative material in the museum’s collection. These two samples were therefore excluded from downstream analyses of endemic diversity.

**Figure 1:**
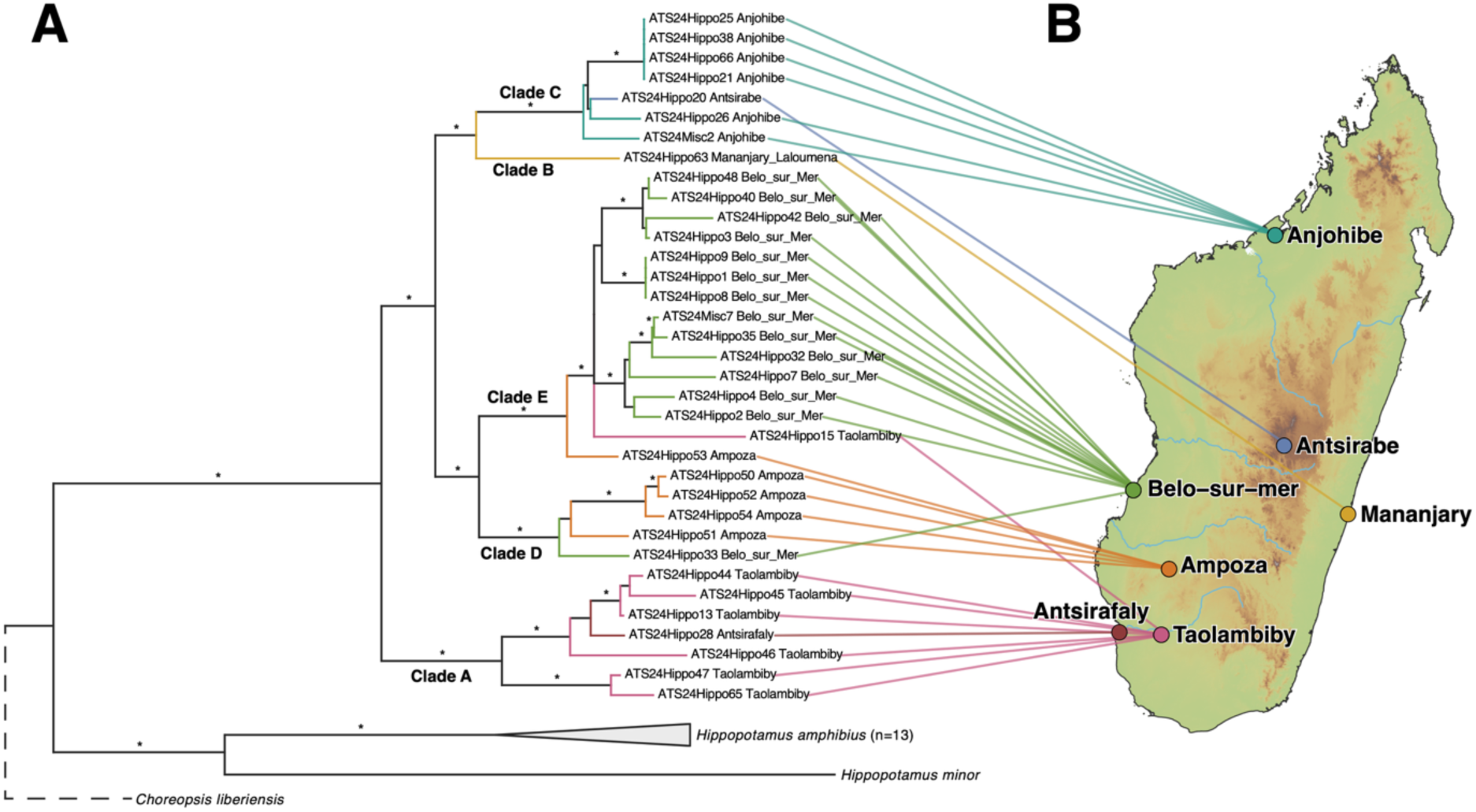
Maximum-likelihood phylogeny of Malagasy hippopotamus mitogenomes with their associated sampling localities in Madagascar. The tree was rooted with the extant pygmy hippopotamus, *Choeropsis liberiensis*. Tips representing newly generated Malagasy specimens are connected to their sampling localities on the map using phytools^94^. Bootstrap support values are shown for major nodes; asterisks indicate nodes with bootstrap support ≥90.

Within the endemic Malagasy *Hippopotamus* clade, we recovered five well-supported deeply divergent lineages (Figures 1 & 2): Clade A: comprised of specimens from southern Madagascar, Taolambiby and Antsirafaly; Clade B: the *H. laloumena* type specimen from the east coast; Clade C: an Antsirabe *H. madagascariensis* from the central highlands and specimens from Anjohibe in northern Madagascar; Clade D: comprised of specimens from southwestern Madagascar, primarily Ampoza but also Belo-sur-Mer; and Clade E: primarily Belo-sur-Mer, also with specimens from Ampoza and Taolambiby (Figure 1 & 2A). Material assigned to *H. lemerlei* spanned Clades A, D and E, rendering it paraphyletic, with *H. madagascariensis* and the larger-bodied *H. laloumena* nested within this diversity. Despite strong association between mtDNA clade and site, several placements departed from this pattern, including an Antsirabe specimen among Anjohibe samples, a Belo-sur-Mer specimen in Clade D, and Taolambiby and Ampoza specimens in Clade E.

### Haplotype structure and mitochondrial diversity

Median-joining network analyses recovered the same major structure observed in the maximum-likelihood trees. In the broader dataset of confirmed endemic Malagasy specimens with >60% mitogenome coverage, 35 sequences collapsed into 17 haplotypes (Figure S3). In the reduced high-coverage dataset, 24 specimens formed 19 haplotypes (Figure 2B). The reduced network more clearly separated the five major Malagasy *Hippopotamus* mtDNA lineages recovered in the phylogenetic analyses.

**Figure 2:**
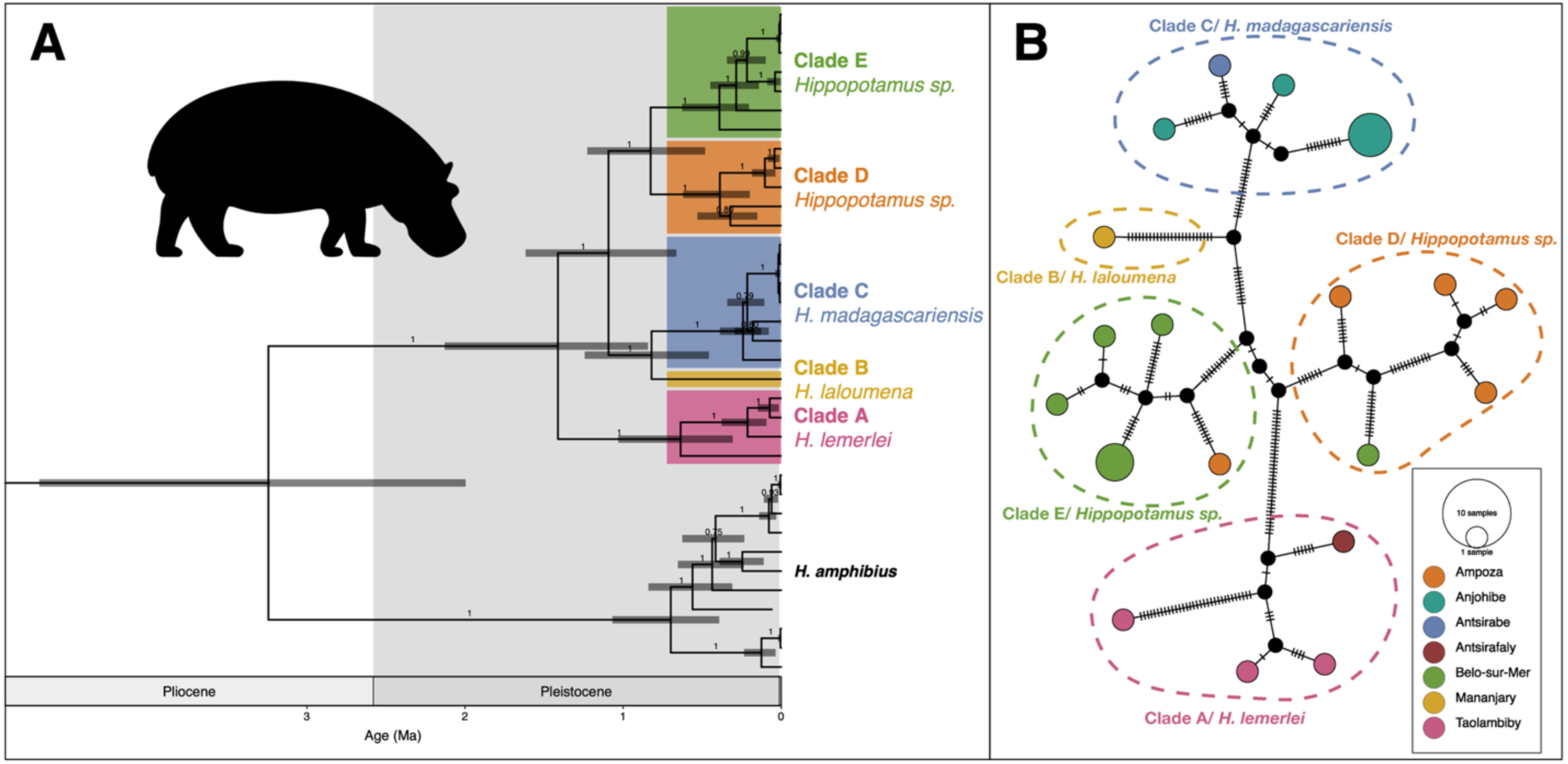
Dated mitochondrial phylogeny and haplotype structure of endemic Malagasy hippopotamuses. (A) Maximum clade credibility tree from the Bayesian tip-dating analysis of the reduced high-coverage dataset, calibrated using the *Choeropsis*–*Hippopotamus* divergence. Node bars show 95% highest posterior density intervals. (B) Median-joining haplotype network for the reduced high-coverage endemic Malagasy dataset. Circle size is proportional to haplotype frequency, colours indicate sampling locality, and hatch marks indicate mutational steps between haplotypes. The two specimens that grouped with modern *H. amphibius* were excluded from the network.

### DNA-only tip-dating analysis

Bayesian tip-dating analyses (Figure 2A) placed the crown age of endemic Malagasy hippopotamuses at 1.41 Ma (95% HPD: 0.90–2.19 Ma), with a divergence from *H. amphibius* lineage at 3.24 Ma (95% HPD: 2.09–4.39 Ma). Within the sampled Malagasy radiation, Clade A diverged from Clades B–E at 1.41 Ma, with crown diversity within Clade A dating to 0.64 Ma. Clades B/C diverged from D/E at 1.09 Ma; B and C split at 0.82 Ma, and D and E at 0.83 Ma. Crown diversity within Clades C–E dates to 0.24–0.39 Ma. These dates were largely consistent, but generally even shallower, when using the divergence between Hippopotamidae and Cetacea as an alternative calibration point (Figure S4).

### Morphological comparisons

Morphological comparisons provided taxonomic context for genetically sampled specimens with associated diagnostic morphological material. The Clade A humerus from Taolambiby fell within the published *H. lemerlei* morphospace (Figure 3C), consistent with the placement of the southern Taolambiby/Antsirafaly lineage as *H. lemerlei*. The Clade E radius from Belo-sur-Mer fell within the *H. lemerlei* morphospace (Figure 3D), providing some postcranial support for a *H. lemerlei*-like morphology in this lineage.

**Figure 3:**
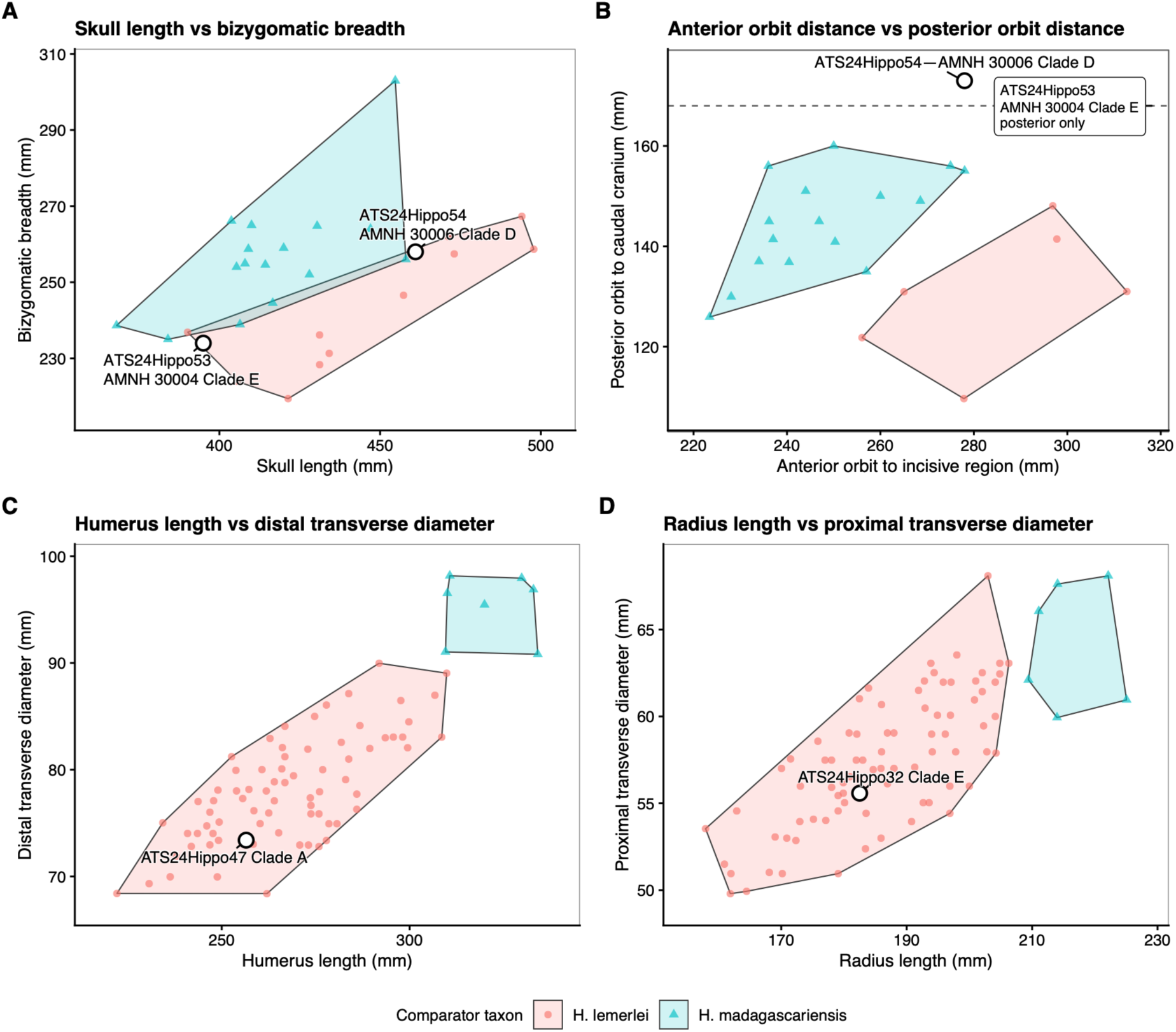
Morphological comparisons of genetically sampled Malagasy hippopotamus specimens compared with published *H. lemerlei* and *H. madagascariensis* specimens. A) Cranial length versus bizygomatic breadth. B) Orbit-position comparison using anterior orbit-to-incisive region distance and posterior orbit-to-caudal cranium distance. C) Humerus length versus distal transverse diameter. D) Radius length versus proximal transverse diameter. Published comparative data were digitized from Stuenes^15^, Fovet et al. ^32^, and Rakotovao et al. ^31^. Polygons represent convex hulls around published specimens assigned to *H. lemerlei* or *H. madagascariensis*. Newly measured specimens were projected onto the plots.

Cranial measurements from the Ampoza specimens representing Clades D and E were more ambiguous. Both crania from the American Museum of Natural History (AMNH) fell within the published Malagasy hippopotamus length–breadth morphospace (Figure 3A), with neither showing the expanded bizygomatic breadth expected for a robust *H. madagascariensis*-like morphology. In contrast, orbit position separated both specimens from the published *H. lemerlei* and *H. madagascariensis* morphospaces (Figure 3B). AMNH FM 30006 (ATS24Hippo54) plotted above both published polygons, while AMNH FM 30004 (ATS24Hippo53), for which only the posterior orbit-to-caudal cranium measurement was available, showed a similarly high value.

Mandibular and dental comparisons were less taxonomically informative. Complete AMNH mandibles were broadly compatible with *H. lemerlei*-like proportions but overlapped substantially with published *H. madagascariensis* ranges (Figure S5). Isolated teeth also fell within overlapping ranges and were not considered diagnostic.

Overall, available morphology supports Clade A as *H. lemerlei*-like and provides some postcranial support for a similar morphology in Clade E, whereas cranial evidence from Clades D and E does not clearly match either published morphospace. Clades B and C were not included in the morphometric comparisons because the sequenced material was too fragmentary to preserve the diagnostic measurements used here. However, Clade B is represented by the *H. laloumena* type specimen, while the Clade C specimen from Antsirabe derives from the type locality of *H. madagascariensis*.

### Stable isotope comparisons

Using a compiled dataset of 76 Malagasy hippopotamus isotope values, including seven genetically sampled endemic specimens, isotope values did not cluster by major mtDNA lineage but were more strongly associated with locality (Figure S6). Antsirafaly and Taolambiby occupied different regions of isotope space despite both localities being primarily associated with Clade A; the Taolambiby Clade E specimen ATS24Hippo15 lacked isotope data. Antsirabe and Anjohibe, which are grouped in Clade C, also strongly differed in isotope values. Conversely, genetically distinct Ampoza/Belo lineages overlapped: Ampoza specimens from different clades had similar values, as did Belo-sur-Mer specimens. All isotope-sampled specimens were consistent with their recorded collecting localities, thus further supporting their documented provenance.

### Total-evidence tip-dating analysis and ancestral state reconstruction

Bayesian total-evidence tip-dating analysis combining mtDNA and morphological characters recovered Malagasy hippos as a single monophyletic radiation that is sister to a clade of *Hippopotamus* species, including *H. amphibius*, with extinct *Hexaprotodon* species and *C. liberiensis* more distantly related (Figure S7). Relationships among Malagasy lineages and divergence times for the radiation on Madagascar generally matched the mtDNA-only analyses. Ancestral-state reconstruction inferred that the most recent common ancestor of the sampled endemic Malagasy lineages was small-bodied, whereas the preceding node was less certain but favoured a medium-to-large body size. The larger-bodied *H. laloumena* is nested within predominantly small-bodied Malagasy lineages, consistent with a secondary increase (reversal) in body size from the reconstructed small-bodied ancestor.

## DISCUSSION

Ancient mitogenomes reveal that Madagascar’s extinct hippopotamuses formed a young, rapidly diversifying endemic radiation derived from a single colonization that strongly contrasts with the ancient radiations of other Malagasy mammalian lineages. We recovered five major, geographically structured mtDNA lineages that do not map cleanly onto traditional taxonomy: material attributed to *H. lemerlei* spans several divergent clades, while the *H. laloumena* type and *H. madagascariensis*-associated material is nested within this broader diversity. Morphology supports some associations but leaves others unresolved, while isotope variation more closely reflects locality than mtDNA lineage. Together, these results reveal a recent island radiation whose evolutionary diversity is only partly represented by current taxonomy.

Our dated phylogenetic reconstructions indicate that much of Madagascar’s hippopotamid diversity accumulated rapidly and recently within Madagascar (Figure 2A). All sampled Malagasy hippopotamuses form a single clade sister to *H. amphibius* and the extinct small-bodied insular *H. minor* (Figure 1). The endemic Malagasy radiation therefore derives from a single mainland African colonization, contradicting hypotheses that the larger-bodied *H. laloumena* represented a separate, later colonization of the island^16,35,36,42^, and the notion that *H. madagascariensis* is instead more closely related to the extant pygmy hippopotamus, *Choeropsis liberiensis*^33,34^. The closer relationship of the Malagasy radiation to *Hippopotamus*, rather than with the extant pygmy hippopotamus *C. liberiensis*, also indicates that reduced body size in Malagasy hippopotamuses evolved independently from that of *Choeropsis*. Although the exact timing of colonization remains uncertain, our fossil-calibrated tip-dated phylogeny places the stem divergence from *H. amphibius* and its recently extinct relatives, at approximately 3.24 Ma (95% HPD: 2.09–4.39 Ma) and the crown age of the sampled Malagasy diversity at approximately 1.41 Ma (95% HPD: 0.90–2.19 Ma). Treating the stem divergence as an upper bound on colonization suggests that hippopotamuses reached Madagascar no earlier than the Pliocene, followed by Pleistocene diversification and persistence into the Holocene.

The mainland progenitor of the Malagasy radiation remains unknown. In the total-evidence analysis, the Malagasy radiation fell outside a clade containing confidently assigned *Hippopotamus* taxa (*H. amphibius, H. gorgops,* and *H. minor*) and Pliocene–Pleistocene African forms such as *protamphibius, karumensis,* and *aethiopicus*, whose generic placement between *Hippopotamus* and *Hexaprotodon* has historically been debated^43,44^. An extinct, unsampled African lineage allied to either genus therefore remains a plausible source. The close mtDNA relationship to *H. amphibius* and *H. minor* identifies its nearest sampled relatives; *H. amphibius* is the nearest sampled living relative, but not necessarily the lineage that colonized Madagascar.

This recent diversification makes Malagasy hippopotamuses exceptional among Madagascar’s endemic mammalian radiations. The four classic endemic nonvolant mammal radiations are all inferred to have colonized Madagascar much earlier in the Eocene–Miocene (∼60–13 Ma)—lemurs (60–34 Ma^22,45^), tenrecs (56–25 Ma^22,46^), euplerid carnivorans (26–18 Ma^22,23^), and nesomyine rodents (24–13 Ma^22,47^). Although the timing and number of lemur colonisations remain debated, even younger estimates for separate Chiromyiformes and Lemuriformes dispersals (∼17.9 and ∼30.4 Ma, respectively)^48,49^ substantially predate the Malagasy hippopotamus radiation. In this respect, Malagasy hippopotamuses represent the youngest nonvolant endemic mammalian radiation on the island, and an unusual example of recent island diversification in a large semi-aquatic herbivore. This demonstrates that Madagascar’s endemic mammal diversity was not solely built by ancient Eocene–Miocene colonisations, but also through more recent rapid diversifications of large-bodied megafauna. Only some bat lineages show comparably recent colonization, involving much smaller-bodied volant mammals with very different dispersal capacities.^21,33,50^

Our results show that traditional species assignments do not map cleanly onto the recovered mtDNA lineages. *H. laloumena* and *H. madagascariensis* are nested within broader mtDNA diversity traditionally attributed to, or morphologically similar to, *H. lemerlei*, rendering *H. lemerlei* paraphyletic (Figures 1–2). This could reflect taxonomic misassignment from fragmentary morphology, retention of ancestral mtDNA lineages, mitochondrial introgression, or derivation of the common ancestor to the *H. laloumena* + *H. madagascariensis* clade from a geographically structured lineage within what is currently recognized as *H. lemerlei.* Comparable paraphyletic mtDNA patterns occur in other systems where named species, geography, and mtDNA lineages do not map perfectly onto one another^51^. Geographically structured non-monophyly has also repeatedly revealed underestimated species diversity, with multiple evolutionary lineages previously subsumed within a single species.^52–54^ Regardless of mechanism, the current morphology-based taxonomy does not fully capture Malagasy hippopotamid diversity.

The limited morphological evidence available for our genetically sampled dataset supports this interpretation. Clade A—comprised of individuals from Taolambiby and Antsirafaly on the southern coast of Madagascar—falls within the expected morphospace of *H. lemerlei*, with Antsirafaly being close to the type locality of the species, Ambolisatra^18^. Clade B is taxonomically anchored by the *H. laloumena* type specimen^16^. Material from Antsirabe and Anjohibe lacked diagnostic morphology, but Clade C includes a specimen from Antsirabe, the type locality of *H. madagascariensis.*^17^ Additionally, previous morphology-based work has suggested that much of the Anjohibe material is more consistent with *H. madagascariensis.*^55^ Therefore, we interpret Clade C as representing *H. madagascariensis*, extending from the central highlands to northern Madagascar. Species assignment is less clear for Clades D and E, which include individuals largely from Belo-sur-Mer and Ampoza, in southwestern Madagascar. The associated specimens from these two clades lack robust *H. madagascariensis*-like cranial morphology, while orbit position— argued to be one of the most diagnostic features^31^—falls outside the published ranges of both *H. lemerlei* and *H. madagascariensis*. We therefore interpret Clades D and E as distinctive *H. lemerlei*-like lineages that likely represent geographically structured variation or undescribed species diversity in western and southwestern Madagascar.

Stable isotope data from the Ampoza- and Belo-sur-Mer-associated lineages (Clades D and E) indicate that mtDNA diversity was not simply structured by dietary differentiation. Isotope values were more strongly associated with locality than mtDNA lineage: genetically distinct Ampoza specimens had very similar isotope signals, while two Belo-sur-Mer specimens from different lineages were similarly congruent. This accords with previous evidence for ecological flexibility and strong geographic influences in Malagasy hippopotamids^37–41,56^. The lack of congruence between genetic lineages and isotope signatures suggests that diversification may have been shaped primarily by geography, demographic history, and local adaptation rather than simple dietary partitioning. There is also the potential for sympatry of multiple clades at a single site, if they co-occurred there at the same time: Belo-sur-Mer (Clades D and E) and Taolambiby (Clades A and E).

The geographic structure of these mtDNA lineages may reflect changing population connectivity among Madagascar’s river systems. The phylogeographic distributions are broadly consistent with the watershed model proposed for Malagasy microendemism, in which low-elevation catchments promote geographic isolation while major drainages originating at higher elevations provide corridors for retreat and dispersal during climatic fluctuations^57^. Such a mechanism may be particularly relevant to hippopotamuses, whose dispersal is closely associated with aquatic habitats, potentially restricting movement across drainage divides while facilitating connectivity along larger river systems. However, the dispersal events we document within Clade D (between Belo-sur-Mer and Ampoza), and Clade E (between Belo-sur-Mer and Taolambiby) traverse between low elevation river drainages, thus arguing against a strict watershed speciation model. Clade C, spanning Antsirabe in the Central Highlands and Anjohibe in northern Madagascar, may represent an even more complex history of connectivity through highland-associated drainage systems. Comparable patterns occur in other Malagasy vertebrates: Central Highland regions have alternately acted as barriers and corridors through time^58–60^, while Pliocene–Pleistocene diversification in geographically structured warty chameleon (*Furcifer verrucosus*) lineages overlaps broadly with the timescale inferred for diversification of Malagasy hippopotamuses^61^.

The survival of multiple lineages until the late Holocene reframes the evolutionary significance of hippopotamid extinction on Madagascar. Rather than representing a minor radiation of peripheral island dwarfs, Malagasy hippopotamuses comprised one of the richest late-surviving assemblages of hippopotamid diversity. Malagasy hippopotamuses retained unusually deep mtDNA structure, whereas *H. amphibius* shows comparatively shallow mitochondrial diversity and limited broad-scale phylogeographic structure^62,63^. Ancient mitogenomes further indicate that sampled Late Pleistocene European hippopotamuses fall within the *H. amphibius* lineage^64^, consistent with a geographically widespread species extending from Africa into Europe. In contrast, Madagascar supported several deeply divergent endemic mtDNA lineages within a far smaller geographic area. Extant *H. amphibius* show strong residency and fidelity to aquatic refugia, with frequent locality-specific mtDNA haplotypes^62^, while large-scale dispersal has been documented in subadult males^65^. In Madagascar, if long-distance dispersal was predominantly male-mediated, such fidelity to aquatic refugia could have promoted persistent female-mediated isolation among drainage systems, allowing deeply divergent, maternally inherited mtDNA lineages to accumulate and persist within a relatively small geographic area.

Although only two hippopotamid species survive today, diversity was considerably greater in the recent past. As hippopotamids dispersed beyond Africa, insular pygmy forms evolved independently on Mediterranean islands including Cyprus, Sicily, Crete and Malta^66–68^, providing repeated, independent examples of island dwarfism and endemism. Most of this diversity, however, was partitioned among separate islands. In contrast, Madagascar supported multiple deeply divergent hippopotamus lineages on a single large island until the late Holocene. Madagascar’s greater area, drainage complexity and habitat heterogeneity may have permitted regional population structure and diversification within a single island, rather than repeated isolation among separate islands.

Body size varied substantially within the Malagasy radiation. *H. lemerlei* and *H. madagascariensis* were both markedly smaller than mainland *H. amphibius*^15^, whereas *H. laloumena* approached mainland body dimensions^14,16^. Ancestral-state reconstruction strongly supported a small-bodied Malagasy crown ancestor (Figure S7), while the immediately preceding node favoured a medium-to-large body size. This implies a transition to reduced size before diversification of the endemic radiation, independently of other dwarf hippopotamids in Europe and mainland Africa. The reconstruction cannot determine whether this change occurred before or after colonization of Madagascar, although an insular reduction following colonization represents the more plausible biogeographic scenario. The nested position of the larger-bodied *H. laloumena* is therefore consistent with a secondary increase in body size within Madagascar.

By the Holocene, much of the Pleistocene hippopotamid diversity of Europe, Asia and Africa had disappeared^64,66,67^, while Madagascar retained at least three named species and several deeply divergent endemic mtDNA lineages. Their extinction within the last millennium therefore erased a substantial component of hippopotamid evolutionary diversity. Ancient mitogenomes, combined with morphology, reveal that this loss encompassed not only the three described species, but also a geographically structured island radiation whose diversity was only partly captured by traditional taxonomy.

## MATERIALS AND METHODS

### Sampling

Analyses were performed on 58 subfossil bone and tooth specimens putatively identified as *Hippopotamus* from a range of sites across Madagascar (Table S1). Sampling included material assigned or tentatively attributed to all three described Malagasy hippopotamus species, *H. lemerlei*, *H. madagascariensis*, and *H. laloumena*. Five samples were newly radiocarbon dated: three at the Center for Accelerator Mass Spectrometry (CAMS), Lawrence Livermore National Laboratory, and two at Beta Analytic. These were combined with eight radiocarbon dates previously generated from specimens included in this study, giving a total of 13 directly dated samples encompassing all named species and the majority of the geographic spread (Table S1). All radiocarbon dates were calibrated with the SHCal20 ^69^ curve using OxCal 4.4 ^70^.

All pre-PCR steps were conducted in a purpose-built ancient DNA laboratory at the American Museum of Natural History, spatially separated and physically isolated from any other molecular laboratories. Strict protocols were followed to minimize contamination of samples with exogenous DNA^71^. These included standard ancient DNA precautions such as the use of full protective clothing, frequent glove changes, routine bleach cleaning and UV irradiation of work surfaces and equipment. These precautions also included the inclusion of negative controls for both DNA extraction and PCR setup. PCR amplification and all downstream procedures were carried out in physically separated DNA laboratories.

### DNA extraction

Surface contamination of each sample was reduced by UV irradiation followed by abrading of the exterior surface of the bone using a Dremel drill with a disposable carborundum disc where appropriate. The sample was then pulverized and 40-100mg of powder used for subsequent DNA extraction. DNA was extracted as per Dabney and Meyer ^72^. This involved an initial digestion in 1mL of 0.5 M EDTA for 60 min, followed by an overnight incubation in 970uL of EDTA and 30 μL of 20 mg/ml proteinase K at 55 °C at 800rpm on a ThermoShaker. Samples were then centrifuged and the supernatant mixed with 13 mL of binding buffer (12.6 mL of Qiagen™ PB buffer, 6.5 μL of Tween-20, and 390 μL of sodium acetate). The mixture was passed through a Qiagen™ MinElute column using a Zymo 15 mL reservoir, washed twice with 700 μL Qiagen™ PE buffer, dried by centrifugation, and eluted twice in 30 μL TE buffer.

### Single-stranded libraries

Single-stranded DNA libraries were prepared using the Santa Cruz Reaction protocol of Kapp, et al. ^73^. Briefly, 20 μL of DNA extract was used as input, and extract concentration quantified using a dsDNA High Sensitivity assay Qubit® (Invitrogen, Waltham, MA) to determine appropriate adapter and single-stranded binding protein concentrations following the protocol’s input-tiering system described by Kapp, et al. ^73^. Extracts exceeding the highest input tier were diluted with ddH2O so that no more than 150 ng DNA was added to the reaction. DNA was first denatured and single-stranded binding proteins annealed by incubation at 95 °C for 3 min followed immediately by cooling on ice for 2 min. Adapter ligation was then carried out using the tier-appropriate adapter dilution together with 26 μL of SCR master mix consisting of 3.75 μL SCR buffer (666 mM Tris-HCl, 132 mM MgCl₂), 0.5 μL ATP (100 mM), 0.5 μL DTT (1 M), 0.625 μL T4 DNA ligase (2,000,000 U/mL), 0.625 μL T4 polynucleotide kinase (10,000 U/mL), and 20 μL PEG 8000 (50%). Reactions were pulse-vortexed five times, briefly spun down, and this process was repeated twice prior to incubation at 37 °C for 45 min. Libraries were then purified using a Qiagen™ MinElute Reaction Cleanup Kit according to the manufacturer’s instructions and eluted in 50 μL EB containing 0.05% Tween 20.

### Library indexing and sequencing

Libraries were double-indexed and amplified in duplicate 50 μL PCRs containing 1.25 U Pfu Turbo Cx Hotstart DNA polymerase (Agilent Technologies), 1× Pfu Turbo Cx buffer, 0.2 mM dNTPs, 0.1 mg/mL BSA, 0.2 μM of each indexing primer, and 12.5 μL of single-stranded library. Cycling conditions were 95 °C for 2 min; 18 cycles of 95 °C for 30 s, 60 °C for 30 s, and 72 °C for 60 s; followed by 72 °C for 10 min. PCR duplicates were pooled by library, purified with 1.0× AxyPrep™ magnetic beads, eluted in 30 μL H2O, quantified using a Qubit fluorometer with a dsDNA High Sensitivity assay, and visualized on an Agilent 4150 TapeStation using a D1000 ScreenTape.

### Mitochondrial enrichment

Libraries were enriched using a custom mitochondrial bait set designed from Malagasy vertebrate mitochondrial genomes (Table S2) and synthesized by Arbor Biosciences (MI, USA) as biotinylated 80-mer RNA baits. DNA-RNA hybridisation enrichment was performed according to manufacturer’s recommendations (MYbaits protocol v3) with the exception that we used 1.25 μL of baits per reaction, changed the incubation step to 55 °C for 15 hrs followed by 50 °C for 16 hrs, and washed the beads three times with 0.1 x SSC and 0.1% SDS (5 min at 50 °C). Enriched libraries were amplified via a final round of “off-bead” PCR split into 4 replicate PCRs (25 μL) each containing 0.5 U AmpliTaq Gold® (Applied Biosystems) DNA polymerase, 1× Gold PCR buffer, 2.5 mM MgCl2, 1 mM dNTPs, and 0.5 μM each of IS5 and IS6^74^. Cycling conditions were as follows: 94 °C for 6 min; 16 cycles of 94 °C for 30 s, 60 °C for 30 s, 72 °C for 40 s; and 72 °C for 10 min. Replicate PCRs were pooled and purified using 1.0× AxyPrep™ magnetic beads (Axygen™), eluted in 30 μL H2O, and quantified on an Agilent TapeStation 4150 using a High Sensitivity D1000 ScreenTape. Enriched libraries were then pooled and sequenced as 150 bp paired-end reads on an Illumina NovaSeq X Plus platform at Azenta/GENEWIZ.

### Data processing

Sequencing data were processed using the nf-core EAGER pipeline v 2.5.1 ^75^ with NextFlow v 22.10.7 ^76^. In short, adapter sequences were removed and paired end reads collapsed using ADAPTER REMOVAL v2.3.2 ^77^ , trimming bases below a minimum Phred quality score of 20, and discarding reads shorter than 20 bp after clipping. Read quality was visualized before and after adapter trimming using fastQC v0.11.9 (http://www.bioinformatics.babraham.ac.uk/projects/fastqc/) to ensure efficient adapter removal. Processed reads were mapped against the mtDNA reference genome of *H. amphibius* (NC_000889) using BWA v 0.7.17-r1188 aln -l 1024, -n 0.01, -o 2; ^78^. BAM files were then filtered using SAMTOOLS v1.12^79^, removing unmapped reads as well as reads mapped with a mapping quality score (MAPQ) less than 25. PCR duplicates were removed using MARKDUPLICATES from the Picard package v 2.26.0 (http://broadinstitute.github.io/picard/). Damage profiles were generated using DamageProfiler v 0.4.9^80^, showing the typical C–T damage profiles characteristic of aDNA (Figure S1). Mapped reads were visualized in Geneious Prime v2019.2.3 (https://www.geneious.com) and consensus sequences generated using a 75% majority-rule, calling N at sites with depth of coverage <3×.

### Comparative hippopotamid sampling

To place the Malagasy hippopotamus mitogenomes in a broader hippopotamid context, we supplemented our newly generated data with publicly available mtDNA data from extant and extinct hippopotamids. Complete mitogenome sequences available from GenBank were downloaded directly, including public mitogenomes of *Hippopotamus amphibius*, *Choeropsis liberiensis*, and *Balaenoptera musculus*. For comparative datasets available only as raw SRA/ENA reads, reads were downloaded and processed through the same pipeline as our newly generated data.

Raw-read datasets included extant *H. amphibius* samples, *Choeropsis liberiensis*, the Late Pleistocene European *Hippopotamus* specimens NK37, Meng19, and HgimEpp from Arnold, et al. ^64^, and the Cypriot pygmy hippopotamus, *H. minor*, from Psonis, et al. ^66^. These reads were adapter-trimmed, mapped to the *H. amphibius* or *C. liberiensis* mt reference genome depending on source species, filtered by mapping quality, deduplicated, and used to generate consensus sequences using the same thresholds applied to the Malagasy specimens. All public accessions, data sources, and reconstructed comparative sequences are listed in Table S3.

An additional *C. liberiensis* mitochondrial genome (C871509) was obtained from a whole-genome sequencing project of hoofed mammals (in prep). In short, an existing DNA extract from Field Museum of Natural History specimen FMNH 140919, a zoo specimen of *Choeropsis liberiensis*, was provided by the Field Museum. A TruSeq PCR-free library was prepared using approximately 450-bp DNA fragments and sequenced at the New York Genome Center to approximately 30× genomic coverage with 150-bp paired-end reads. Raw-read quality was assessed with FastQC, and reads were trimmed with Trimmomatic^81^. Trimmed reads were assembled into contigs de novo with MEGAHIT^82^ using k-mer sizes from 39 to 95 in increments of eight, with the mercy rescue option disabled. MEC^83^ was used to identify and split putatively misassembled contigs, which were then scaffolded against the *C. liberiensis* reference genome (GCA_023065765) using RagTag^84^. The mitochondrial sequence was identified by searching the resulting assembly with BLAST using the reference mitogenome (NC_020697) as a query; the best match was then selected based on alignment length and sequence identity.

### Mitogenome alignment and dataset construction

Published and reconstructed comparative mtDNA sequences were added to the newly generated Malagasy hippopotamus consensus sequences with >50% mitogenome coverage, and the full dataset was aligned using MUSCLE v3.8.425 ^85^ as implemented in Geneious Prime v2019.2.3. After alignment, four datasets were generated for downstream analyses. Full datasets included all newly generated Malagasy hippopotamus consensus sequences that met the minimum coverage threshold, together with downloaded and reconstructed comparative hippopotamid sequences. Reduced datasets were restricted to specimens with at least 75% of the mitogenome and were used for haplotype-network, and Bayesian tip-dating analyses. For both the full and reduced datasets, we generated one alignment using *Choeropsis liberiensis* as the outgroup and a second alignment including blue whale, *Balaenoptera musculus*, as a more distant cetacean outgroup with the poorly aligned control region excluded.

### Phylogenetic analysis

Each of the four datasets was partitioned independently using PartitionFinder 2.1.1^86^ selecting the best-fitting partitioning scheme and substitution model under the Bayesian information criterion (Table S4).

Maximum-likelihood phylogenetic analyses were conducted on the two full mitogenome alignments using RAxML v8.2.12^87^, applying the PartitionFinder partitioning schemes selected under the Bayesian information criterion (Table S4) and a GTR+G substitution model for all partitions. Node support was assessed using 1,000 rapid bootstrap replicates.

Population structure within Malagasy hippopotamuses was investigated using median-joining haplotype networks constructed in PopART v1.7.1^88^. Haplotype networks were constructed for all confirmed Malagasy specimens with >60% mitogenome coverage and for a reduced dataset of specimens with >75% mitogenome coverage. The two specimens that grouped with modern *H. amphibius* in phylogenetic analyses were excluded from Malagasy haplotype network analyses.

Bayesian tip-dating analyses were performed in BEAST 2.7.7^89^ using BEAUTi to generate XML files. Analyses were run on the two reduced high-coverage mitogenome datasets. The dataset-specific PartitionFinder schemes were implemented in BEAST using substitution models selected under the Bayesian information criterion (Table S4). Site models were unlinked across partitions, while the tree and clock models were shared across partitions. An optimized relaxed molecular clock was used, with a uniform prior on the clock rate from 0 to 1 × 10^−5^ substitutions/site/year, and a Bayesian skyline tree prior used. In the primary analysis, *Choeropsis liberiensis* was used as the outgroup and the analysis was calibrated using a prior on the divergence between *Choeropsis* and *Hippopotamus* (normal prior: mean = 8.1 Ma, SD = 0.31 Ma; following ^66^). In a second analysis *Balaenoptera musculus* was included and a whale–hippopotamid divergence calibration (lognormal prior: offset = 52.4 Ma, M = 1.0, S = 1.0; based on the *Himalayacetus subathuensis* fossil minimum^90,91^) applied to assess sensitivity to deeper outgroup sampling and calibration choice. Radiocarbon-dated specimens were assigned fixed tip ages with the median calibrated radiocarbon date. Undated specimens were assigned uniform tip-date priors based on site-specific chronological context (Table S5). We ran three independent MCMC chains for 50 million steps, sampling every 5000 steps. Convergence and effective sample sizes were assessed in Tracer v1.7.2^92^. Runs were combined in LogCombiner after discarding the first 10% as burn-in, and maximum clade credibility trees were generated in TreeAnnotator using median node ages.

### Morphological measurements

To compare genetically sampled specimens with published morphological assignments, we measured available cranial, mandibular, dental, and postcranial material following Stuenes ^15^, Fovet, et al. ^32^, and Rakotovao, et al. ^31^. As many specimens were incomplete, we focused only on measurements that could be confidently matched to published comparative data. For crania, we compared AMNH specimens from Ampoza with published measurements of *H. lemerlei* and *H. madagascariensis* using bivariate plots of cranial length and breadth, upper M3 dimensions, and orbit position. For the orbit-position comparison, we used the distance from the anterior border of the orbit to the anterior incisive region and the distance from the posterior border of the orbit to the caudal cranium/external occipital region^31,32^. For postcranial material, we compared a Taolambiby humerus and a Belo-sur-Mer radius length and transverse diameter with the published morphospaces for *H. lemerlei* and *H. madagascariensis*. Published comparative points were digitized from the original figures and used to draw convex hulls for each species. Newly measured specimens were then plotted onto these published comparative spaces but were not included in the polygons. Mandibular measurements and isolated lower teeth were compared with published *H. lemerlei* and *H. madagascariensis* measurement ranges, rather than used for morphospace plots.

### Stable isotope comparisons

To assess ecological variation among Malagasy hippopotamuses, we compiled published stable carbon (δ¹³C) and nitrogen (δ¹⁵N) isotope data from Crowley, et al. ^39^ and supplemented this dataset with newly generated values from this study. In herbivores, δ¹³C primarily reflects plant-resource use, particularly C3 versus C4/CAM vegetation, while δ¹⁵N can provide additional information on habitat and local environmental conditions. We therefore used these data to assess whether ecological variation corresponded more closely to mtDNA lineage or geography. The comparative isotope dataset was restricted to localities represented in the genetic dataset: Antsirabe, Anjohibe, Antsirafaly, Taolambiby, Ampoza, and Belo-sur-Mer. Original source attributions for published isotope values are detailed in Table S6. Because isotope sampling was uneven across localities and mtDNA lineages, and because several mtDNA clades are represented by only one or a few isotope-sampled specimens, these comparisons were treated as descriptive ecological context rather than as a formal test of dietary differentiation among clades. Isotope values were compared by locality and by mtDNA lineage to assess whether ecological variation more closely tracked geography or genetic structure.

### Total Evidence phylogenetic analysis

To investigate the placement of the Malagasy radiation relative to extinct Hippopotamidae, we assembled a total-evidence dataset combining representative mitochondrial genomes with a 37-character morphological matrix derived from a published hippopotamid dataset^43^. The morphological matrix was expanded through additional scoring of the insular dwarf hippopotamuses, including *H. lemerlei*, *H. madagascariensis* and the Cypriot *H. minor*, using published descriptions, measurements and figures^15,93^. Genetically sampled representatives were used for *H. lemerlei* (ATS24Hippo28, Clade A) and *H. madagascariensis* (ATS24Hippo20, Clade C). Clade B/*H. laloumena* (ATS24Hippo63), Clade D (ATS24Hippo50) and Clade E (ATS24Hippo48) were represented by their mt genomes but morphological characters were coded as missing because diagnostic morphological material could not be confidently associated with the sequenced representatives. The matrix additionally included extant *H. amphibius* and *Choeropsis liberiensis* and a range of extinct African and Asian hippopotamids, including *H. gorgops*, *H. minor* and fossil taxa historically attributed to *Hexaprotodon*.

Morphological characters were partitioned according to the number of character states and analysed under ascertainment-corrected Lewis Mk models. Mitochondrial partitions followed the substitution-model framework used in the mt phylogenetic analyses, with a relaxed molecular clock applied to the mt data and a separate strict clock for morphology. The ages of 11 morphology-only fossil taxa were estimated jointly with the phylogeny using priors based on published stratigraphic estimates (Table S7). Ten were assigned uniform priors spanning their estimated age ranges, while *Saotherium* cf. *mingoz* was assigned a normal prior (mean 5.26 Ma, SD 0.29 Ma). The age of *Hex. bruneti* was fixed at 2.50 Ma. Analyses used a fossilized birth-death tree prior that allowed sampled ancestors in BEAST 2.7.7. The *Choeropsis*-*Hippopotamus* divergence was assigned a normal prior with a mean of 8.1 Ma and SD of 0.31 Ma. Three independent chains were run for 100 million MCMC generations, with convergence and effective sample sizes assessed in Tracer, log and trees files combined in LogCombiner discarding the first 10% as burn-in and a maximum-clade-credibility tree summarized with median heights in TreeAnnotator.

### Ancestral State Reconstruction

To examine body-size evolution within Hippopotamidae, terminal taxa in the total-evidence tree were assigned to broad relative adult body-size categories (small, medium or large) based on published morphological descriptions and size comparisons^14–16,43,44^. These categories were intentionally qualitative rather than fixed body-mass thresholds because reliable numerical mass estimates are unavailable for many fossil taxa. Malagasy Clades A, C, D and E were coded as small, *H. laloumena* (Clade B) as medium.

Ancestral-state reconstruction was performed on the total-evidence maximum clade credibility tree in R using phytools^94^. Five alternative transition models were compared using the sample-size-corrected Akaike information criterion (AICc): an ordered equal-rates model, an ordered model with separate rates for increases and decreases, and unordered equal-rates (ER), symmetric (SYM) and all-rates-different (ARD) models. To account for uncertainty in the transition model, ancestral-state probabilities were averaged across models according to their AICc weights. Equal prior probabilities were assigned to the three states at the root. Because fossilized birth-death trees contain zero-length branches associated with sampled ancestors, these branches were retained rather than artificially extended.

## ACKNOWLEDGEMENTS

We would like to thank Jin Meng, Judith Galkin and Alana Gishlick of the American Museum of Natural History Division of Paleontology for assistance with specimen access and sampling, and Ursula B. Göhlich and the Natural History Museum Vienna for access to *Hippopotamus* specimens in their collection. We thank FMNH Mammalogy, Bill Stanley and Adam Ferguson for access to pygmy hippopotamus tissues and provision of the DNA extract. We thank Lauren Clark, Seth Cunningham, Melina Giakoumis, Mohammad Faiz and Cheryl Y. Hayashi for their support of our work in the American Museum of Natural History Ancient Biomolecular Laboratory (AbLab). We are grateful to faculty and students from the Université d’Antananarivo and Fordham University for assistance with fieldwork. We thank the Ministère des Mines for issuing excavation and export permits, the Direction Régionale Atsimo-Andrefana for facilitating excavation, and the Ministry of Culture and Communication, particularly its Direction du Patrimoine, for assistance with the excavation. Material used in this study was collected with support from grants from the National Science Foundation, National Geographic Society, National Oceanic and Atmospheric Administration, and Smithsonian Institution. This research was funded by the U.S. National Science Foundation (DBI-2029955 awarded to C.J.R. and B.T.S.; RAPID DEB-1931213 awarded to E.H. and C.J.R.) and the Frick Fund, AMNH Division of Paleontology. Laboratory work in the American Museum of Natural History Ancient Biomolecular Laboratory was supported by the U.S. National Science Foundation (DBI-2015389). This work used Expanse at the San Diego Supercomputer Center through allocation BIO240142 from the Advanced Cyberinfrastructure Coordination Ecosystem: Services & Support (ACCESS) program^95^, which is supported by U.S. National Science Foundation grants #2138259, #2138286, #2138307, #2137603, and #2138296.

## DATA AVAILABILITY STATEMENT

The mitochondrial genome sequences generated in this study will be made publicly available in GenBank upon publication of the peer-reviewed article. In the interim, data may be made available from the corresponding author upon reasonable request.

## REFERENCES

1. Burney, D.A. et al. A chronology for late prehistoric Madagascar. J. Hum. Evol. 47, 25–63 (2004).

2. Lewis, M.E. et al. New large species of *Cryptoprocta* (Carnivora, Eupleridae) and new specimens of *C. Spelea* from the Holocene of Tsimanampesotse National Park, southwest Madagascar. J. Vertebr. Paleontol., e2698802 (2026).

3. Godfrey, L.R. et al. Patterns of late Holocene and historical extinctions on Madagascar. Camb. Prism. Extinct. 3, e9 (2025).

4. Crowley, B.E. A refined chronology of prehistoric Madagascar and the demise of the megafauna. Quat. Sci. Rev. 29, 2591–2603 (2010).

5. Antonelli, A. et al. Madagascar’s extraordinary biodiversity: Evolution, distribution, and use. Science 378, eabf0869 (2022).

6. Hekkala, E. et al. Paleogenomics illuminates the evolutionary history of the extinct Holocene ’horned’ crocodile of Madagascar, Voay robustus. Commun. Biol. 4, 505 (2021).

7. Grealy, A. et al. Molecular exploration of fossil eggshell uncovers hidden lineage of giant extinct bird. Nat. Commun. 14, 914 (2023).

8. Marciniak, S., et al. Evolutionary and phylogenetic insights from a nuclear genome sequence of the extinct, giant, “subfossil” koala lemur *Megaladapis edwardsi*. Proc. Natl. Acad. Sci. U.S.A. 118(2021).

9. Kistler, L. et al. Comparative and population mitogenomic analyses of Madagascar’s extinct, giant ‘subfossil’ lemurs. J. Hum. Evol. 79, 45–54 (2015).

10. Kehlmaier, C. et al. Ancient DNA elucidates the lost world of western Indian Ocean giant tortoises and reveals a new extinct species from Madagascar. Sci. Adv. 9, eabq2574 (2023).

11. Hempel, E. et al. Blue turns to gray: Paleogenomic insights into the evolutionary history and extinction of the blue antelope (*Hippotragus leucophaeus*). Mol. Biol. Evol. 39(2022).

12. Mitchell, K.J. & Rawlence, N.J. Examining natural history through the lens of palaeogenomics. Trends Ecol Evol 36, 258–267 (2021).

13. Thomson, V.A., et al. Using ancient DNA to study the origins and dispersal of ancestral Polynesian chickens across the Pacific. Proc. Natl. Acad. Sci. U.S.A. 111, 4826–31 (2014).

14. Faure, M., Guerin, C., Genty, D., Gommery, D. & Ramanivosoa, B. Le plus ancien hippopotame fossile (*Hippopotamus laloumena*) de Madagascar (Belobaka, Province de Mahajanga). C. R. Palevol 9, 155–162 (2010).

15. Stuenes, S. Taxonomy, habits, and relationships of the subfossil Madagascan hippopotami *Hippopotamus lemerlei* and *H. madagascariensis*. J. Vertebr. Paleontol. 9, 241–268 (1989).

16. Faure, M. & Guerin, C. Hippopotamus laloumena nov. sp., la troisieme espece d’hippopotame holocene de Madagascar. C. R. Acad. Sci. Ser. II 310, 1299–1305 (1990).

17. Guldberg, G. Undersøgelser over en subfossil flodhest fra Madagascar. Christiana Videnskabs-Selskab Forhandlinger 6, 1–24 (1883).

18. Grandidier, A. Sur les découvertes zoologiques faites récemmment à Madagascar. Annales des Sciences naturelles 5 Zoologie et Paléontologie 10, 375–378 (1868).

19. Yoder, A.D. & Nowak, M.D. Has vicariance or dispersal been the predominant biogeographic force in Madagascar? Only time will tell. Annu. Rev. Ecol. Evol. Syst. 37, 405–431 (2006).

20. Ali, J.R. & Vences, M. Mammals and long-distance over-water colonization: The case for rafting dispersal; the case against phantom causeways. J. Biogeogr. 46, 2632–2636 (2019).

21. Samonds, K.E., et al. Spatial and temporal arrival patterns of Madagascar’s vertebrate fauna explained by distance, ocean currents, and ancestor type. Proc. Natl. Acad. Sci. U.S.A. 109, 5352–7 (2012).

22. Poux, C. et al. Asynchronous colonization of Madagascar by the four endemic clades of primates, tenrecs, carnivores, and rodents as inferred from nuclear genes. Syst. Biol. 54, 719–30 (2005).

23. Yoder, A.D. et al. Single origin of Malagasy Carnivora from an African ancestor. Nature 421, 734–7 (2003).

24. Yoder, A.D., Cartmill, M., Ruvolo, M., Smith, K. & Vilgalys, R. Ancient single origin for Malagasy primates. Proc. Natl. Acad. Sci. U.S.A. 93, 5122–6 (1996).

25. Douady, C.J., Catzeflis, F., Kao, D.J., Springer, M.S. & Stanhope, M.J. Molecular evidence for the monophyly of Tenrecidae (Mammalia) and the timing of the colonization of Madagascar by Malagasy tenrecs. Mol. Phylogenet. Evol. 22, 357–63 (2002).

26. Masters, J.C. et al. Biogeographic mechanisms involved in the colonization of Madagascar by African vertebrates: Rifting, rafting and runways. J. Biogeogr. 48, 492–510 (2021).

27. Ali, J.R. & Hedges, S.B. A review of geological evidence bearing on proposed Cenozoic land connections between Madagascar and Africa and its relevance to biogeography. Earth-Sci. Rev. 232, 104103 (2022).

28. Mazza, P.P.A. If hippopotamuses cannot swim, how did they colonize islands? Lethaia 47, 494–499 (2014).

29. van der Geer, A.A.E., Anastasakis, G. & Lyras, G.A. If hippopotamuses cannot swim, how did they colonize islands: a reply to Mazza. Lethaia 48, 147–150 (2014).

30. Godfrey, L.R. et al. Teasing apart impacts of human activity and regional drought on Madagascar’s large vertebrate fauna: insights from new excavations at Tsimanampesotse and Antsirafaly. Front. Ecol. Evol. 9, 742203 (2021).

31. Rakotovao, M., Lignereux, Y., Orliac, M.J., Duranthon, F. & Antoine, P.-O. *Hippopotamus lemerlei* Grandidier, 1868 et *Hippopotamus madagascariensis* Guldberg, 1883 (Mammalia, Hippopotamidae) : anatomie cranio-dentaire et revision systematique. Geodiversitas 36, 117–161 (2014).

32. Fovet, W., Faure, M. & Guerin, C. *Hippopotamus guldbergi* n. sp.: revision du statut d’*Hippopotamus madagascariensis* Guldberg, 1883, apres plus d’un siecle de malentendus et de confusions taxonomiques. Zoosystema 33, 61–82 (2011).

33. Samonds, K.E. et al. Imperfect isolation: factors and filters shaping Madagascar’s extant vertebrate fauna. PLOS ONE 8, e62086 (2013).

34. Harris, J.M. Family Hippopotamidae. in Koobifora Research Project (ed. Harris, J.M.) 31–85 (Oxford University Press, Oxford, 1990).

35. Weston, E.M. & Lister, A.M. Insular dwarfism in hippos and a model for brain size reduction in *Homo floresiensis*. Nature 459, 85–8 (2009).

36. Stankiewicz, J., Thiart, C., Masters, J.C. & de Wit, M.J. Did lemurs have sweepstake tickets? An exploration of Simpson’s model for the colonization of Madagascar by mammals. J. Biogeogr. 33, 221–235 (2006).

37. Hixon, S. Mid-Holocene aridity recorded in pygmy hippo and giant tortoise bone from southwest Madagascar. Proc. R. Soc. B 292, 20250493 (2025).

38. Hansford, J.P. & Turvey, S.T. Dietary isotopes of Madagascar’s extinct megafauna reveal holocene browsing and grazing guilds. Biol. Lett. 18, 20220094 (2022).

39. Crowley, B.E., Godfrey, L.R. & Samonds, K.E. What can hippopotamus isotopes tell us about past distributions of C4 grassy biomes on Madagascar? Plants People Planet 5, 997–1010 (2023).

40. Hixon, S.W. et al. Late Holocene spread of pastoralism coincides with endemic megafaunal extinction on Madagascar. Proc. R. Soc. B 288, 20211204 (2021).

41. Godfrey, L.R. & Crowley, B.E. Madagascar’s ephemeral palaeo-grazer guild: who ate the ancient C4 grasses? Proc. R. Soc. B 283(2016).

42. Monnier, L. & Lamberton, C. Note sur des ossements subfossiles de la région de Mananjary. *Bulletin de l’Académie malgache*, Nouvelle Série 3, 211–213 (1922).

43. Boisserie, J.-R., Likius, A., Vignaud, P. & Brunet, M. A new Late Miocene hippopotamid from Toros-Menalla, Chad. J. Vertebr. Paleontol. 25, 665–673 (2005).

44. Boisserie, J.R. Hippopotamidae (Cetartiodactyla, Hippopotamoidea) from Kanapoi, Kenya, and the taxonomic status of the late early Pliocene hippopotamids from the Turkana Basin. J. Hum. Evol. 140, 102377 (2020).

45. Everson, K.M. et al. Multiple bursts of speciation in Madagascar’s endangered lemurs. Nat. Commun. 16, 1–15 (2025).

46. Everson, K.M., Soarimalala, V., Goodman, S.M. & Olson, L.E. Multiple loci and complete taxonomic sampling resolve the phylogeny and biogeographic history of tenrecs (Mammalia: Tenrecidae) and reveal higher speciation rates in Madagascar’s humid forests. Syst. Biol. 65, 890–909 (2016).

47. Steppan, S.J. & Schenk, J.J. Muroid rodent phylogenetics: 900-species tree reveals increasing diversification rates. PLOS ONE 12, e0183070 (2017).

48. Gunnell, G.F. et al. Fossil lemurs from Egypt and Kenya suggest an African origin for Madagascar’s aye-aye. Nat. Commun. 9, 3193 (2018).

49. Ali, J.R. & Hedges, S.B. The colonisation of Madagascar by land-bound vertebrates. Biol. Rev. Camb. Philos. Soc. 98, 1583–1606 (2023).

50. Christidis, L., Goodman, S.M., Naughton, K. & Appleton, B. Insights into the evolution of a cryptic radiation of bats: dispersal and ecological radiation of Malagasy Miniopterus (Chiroptera: Miniopteridae). PLOS ONE 9, e92440 (2014).

51. Funk, D.J. & Omland, K.E. Species-level paraphyly and polyphyly: frequency, causes, and consequences, with insights from animal mitochondrial DNA. Annu. Rev. Ecol. Evol. Syst. 34, 397–423 (2003).

52. Hekkala, E. et al. An ancient icon reveals new mysteries: mummy DNA resurrects a cryptic species within the Nile crocodile. Mol. Ecol. 20, 4199–4215 (2011).

53. Wan, T., He, K. & Jiang, X.-L. Multilocus phylogeny and cryptic diversity in Asian shrew-like moles (*Uropsilus*, Talpidae): implications for taxonomy and conservation. BMC Evol. Biol. 13, 1–13 (2013).

54. Caminer, M.A. et al. Systematics of the *Dendropsophus leucophyllatus* species complex (Anura: Hylidae): Cryptic diversity and the description of two new species. PLOS ONE 12, e0171785 (2017).

55. Shackelton, A. Master’s Thesis, Northern Illinois University (2021).

56. Joseph, G.S. et al. Resolving the biome debate: ecological and palaeo-data concur that the extensive tree-limited grassy biomes of central and north-western Madagascar are largely anthropogenic. Biol. Rev. Camb. Philos. Soc. (2026).

57. Wilme, L., Goodman, S.M. & Ganzhorn, J.U. Biogeographic evolution of Madagascar’s microendemic biota. Science 312, 1063–5 (2006).

58. Florio, A.M., Ingram, C.M., Rakotondravony, H.A., Louis, E.E. & Raxworthy, C.J. Detecting cryptic speciation in the widespread and morphologically conservative carpet chameleon (*Furcifer lateralis*) of Madagascar. J. Evol. Biol. 25, 1399–414 (2012).

59. Yoder, A.D., et al. Geogenetic patterns in mouse lemurs (genus *Microcebus*) reveal the ghosts of Madagascar’s forests past. Proc. Natl. Acad. Sci. U.S.A. 113, 8049–56 (2016).

60. Everson, K.M., Jansa, S.A., Goodman, S.M. & Olson, L.E. Montane regions shape patterns of diversification in small mammals and reptiles from Madagascar’s moist evergreen forest. J. Biogeogr. 47, 2059–2072 (2020).

61. Florio, A.M. & Raxworthy, C.J. A phylogeographic assessment of the Malagasy giant chameleons (*Furcifer verrucosus* and *Furcifer oustaleti*). PLOS ONE 11, e0154144 (2016).

62. Okello, J.B.A., Nyakaana, S., Masembe, C., Siegismund, H.R. & Arctander, P. Mitochondrial DNA variation of the common hippopotamus: evidence for a recent population expansion. Heredity 95, 206–215 (2005).

63. Stoffel, C. et al. Genetic consequences of population expansions and contractions in the common hippopotamus (*Hippopotamus amphibius*) since the Late Pleistocene. Mol. Ecol. 24, 2507–20 (2015).

64. Arnold, P. et al. Ancient DNA and dating evidence for the dispersal of hippos into central Europe during the last glacial. Curr. Biol. 35, 5363–5371 (2025).

65. Stears, K., Nunez, T.A., Muse, E.A., Mutayoba, B.M. & McCauley, D.J. Spatial ecology of male hippopotamus in a changing watershed. Sci. Rep. 9, 15392 (2019).

66. Psonis, N. et al. Mitochondrial sequences of the extinct Cypriot pygmy hippopotamus confirm its phylogenetic placement. Zool. J. Linn. Soc. 196, 979–989 (2022).

67. Marra, A.C. Pleistocene hippopotamuses of Mediterranean islands: looking for ancestors. in Proceedings of the International Symposium “Insular Vertebrate Evolution: the Palaeontological Approach” Vol. 12 193–204 (Societat d’Història Natural de Balears, Mallorca, Spain, 2005).

68. Lomolino, M.V. et al. Of mice and mammoths: generality and antiquity of the island rule. J. Biogeogr. 40, 1427–1439 (2013).

69. Hogg, A.G. et al. SHCal20 Southern Hemisphere Calibration, 0–55,000 Years cal BP. Radiocarbon 62, 759–778 (2020).

70. Ramsey, C.B. Bayesian analysis of radiocarbon dates. Radiocarbon 51, 337–360 (2009).

71. Fulton, T.L. Setting up an ancient DNA laboratory. Methods Mol. Biol. 840, 1–11 (2012).

72. Dabney, J. & Meyer, M. Extraction of highly degraded DNA from ancient bones and teeth. Methods Mol. Biol. 1963, 25–29 (2019).

73. Kapp, J.D., Green, R.E. & Shapiro, B. A fast and efficient single-stranded genomic library preparation method optimized for ancient DNA. J. Hered. 112, 241–249 (2021).

74. Meyer, M. & Kircher, M. Illumina sequencing library preparation for highly multiplexed target capture and sequencing. Cold Spring Harb. Protoc. 2010, pdb prot5448 (2010).

75. Fellows Yates, J.A., et al. Reproducible, portable, and efficient ancient genome reconstruction with nf-core/eager. Peerj 9, e10947 (2021).

76. Di Tommaso, P. et al. Nextflow enables reproducible computational workflows. Nat. Biotechnol. 35, 316–319 (2017).

77. Schubert, M., Lindgreen, S. & Orlando, L. AdapterRemoval v2: rapid adapter trimming, identification, and read merging. BMC Res. Notes 9, 88 (2016).

78. Li, H. & Durbin, R. Fast and accurate short read alignment with Burrows-Wheeler transform. Bioinformatics 25, 1754–60 (2009).

79. Li, H. et al. The sequence alignment/map format and SAMtools. Bioinformatics 25, 2078–9 (2009).

80. Neukamm, J., Peltzer, A. & Nieselt, K. DamageProfiler: fast damage pattern calculation for ancient DNA. Bioinformatics 37, 3652–3653 (2021).

81. Bolger, A.M., Lohse, M. & Usadel, B. Trimmomatic: a flexible trimmer for Illumina sequence data. Bioinformatics 30, 2114–20 (2014).

82. Li, D., Liu, C.M., Luo, R., Sadakane, K. & Lam, T.W. MEGAHIT: an ultra-fast single-node solution for large and complex metagenomics assembly via succinct de Bruijn graph. Bioinformatics 31, 1674–6 (2015).

83. Wu, B. et al. MEC: Misassembly Error Correction in Contigs based on Distribution of Paired-End Reads and Statistics of GC-contents. IEEE Trans. Comput. Biol. Bioinform. 17, 847–857 (2020).

84. Alonge, M. et al. Automated assembly scaffolding using RagTag elevates a new tomato system for high-throughput genome editing. Genome Biol. 23, 258 (2022).

85. Edgar, R.C. MUSCLE: multiple sequence alignment with high accuracy and high throughput. Nucleic Acids Res 32, 1792–7 (2004).

86. Lanfear, R., Frandsen, P.B., Wright, A.M., Senfeld, T. & Calcott, B. PartitionFinder 2: new methods for selecting partitioned models of evolution for molecular and morphological phylogenetic analyses. Mol. Biol. Evol. 34, 772–773 (2016).

87. Stamatakis, A. RAxML version 8: a tool for phylogenetic analysis and post-analysis of large phylogenies. Bioinformatics 30, 1312–1313 (2014).

88. Leigh, J.W. & Bryant, D. POPART: full-feature software for haplotype network construction. Methods Ecol. Evol. 6, 1110–1116 (2015).

89. Bouckaert, R. et al. BEAST 2.5: An advanced software platform for Bayesian evolutionary analysis. PLOS Comput. Biol. 15, e1006650 (2019).

90. Bajpai, S. & Gingerich, P.D. A new Eocene archaeocete (Mammalia, Cetacea) from India and the time of origin of whales. Proc. Natl. Acad. Sci. U.S.A. 95, 15464–8 (1998).

91. Benton, M.J. et al. Constraints on the timescale of animal evolutionary history. *Palaeontol*. Electron. 18, 1FC (2015).

92. Rambaut, A., Drummond, A.J., Xie, D., Baele, G. & Suchard, M.A. Posterior summarization in Bayesian phylogenetics using Tracer 1.7. Syst. Biol. 67, 901–904 (2018).

93. Lyras, G.A., van der Geer, B.G.M., Kafousia, M.S. & van der Geer, A.A.E. On the cranial anatomy of the smallest insular hippopotamus (Cyprus, Pleistocene). Boll. Soc. Paleontol. Ital. 64, 337–348 (2025).

94. Revell, L.J. phytools 2.0: an updated R ecosystem for phylogenetic comparative methods (and other things). Peerj 12, e16505 (2024).

95. Boerner, T.J., Deems, S., Furlani, T.R., Knuth, S.L. & Towns, J. ACCESS: advancing innovation: NSF’s advanced cyberinfrastructure coordination ecosystem: services & support. in Practice and Experience in Advanced Research Computing 2023: Computing for the Common Good 173–176 (Association for Computing Machinery, Portland, OR, USA, 2023).

