## Supplementary Material for "Ancient mitogenomes reveal unexpected diversity in Madagascar’s extinct hippopotamuses"

**This PDF file includes:**

Tables S1 to S7
Figures S1 to S7
Supplementary References

**Table S1: Malagasy hippopotamus specimens included in this study.** Specimen identifiers, museum and field information, sampled skeletal element, locality, and available radiocarbon dates. Calibrated ages are reported as median calibrated age and 95.4% probability intervals. Identifiers reproduce field, excavation, or specimen labels associated with the material and do not necessarily represent formal field numbers. Radiocarbon ages and their uncertainties are reported in $^{14}$C years BP, and calibrated ages in cal BP.

| ATS Sample # | Museum accession | Identifier | Element | Locality | Date number | Carbon Date | Uncertainty | Calibrated Median | Calibrated Upper %95.4 | Calibrated Lower %95.4 | Date Reference |
|---|---|---|---|---|---|---|---|---|---|---|---|
| **ATS24Hippo1** | AMNH uncatalogued | SQ 3-5 100+ cm | Rib | Belo-sur-Mer | | | | | | | |
| **ATS24Hippo2** | AMNH uncatalogued | SQ 1Layer 6 60-70cm BD | Cervical vertebra | Belo-sur-Mer | | | | | | | |
| **ATS24Hippo3** | AMNH uncatalogued | SQ 1 Layer 0-20cm | Molar | Belo-sur-Mer | | | | | | | |
| **ATS24Hippo4** | AMNH uncatalogued | SQ 1 Layer 0-20cm | Proximal rib | Belo-sur-Mer | CAMS 194198 | 2455 | 40 | 2469 | 2703 | 2346 | this paper |
| **ATS24Hippo5** | AMNH uncatalogued | Slump | Molar | Belo-sur-Mer | | | | | | | |
| **ATS24Hippo6** | AMNH uncatalogued | AZ1.1 | Vertebra | Belo-sur-Mer | | | | | | | |
| **ATS24Hippo7** | AMNH uncatalogued | AZ0.8 | Tooth | Belo-sur-Mer | | | | | | | |
| **ATS24Hippo8** | AMNH uncatalogued | SQ 2-3-4 Oh slump | Premolar and carpal and miscellaneous | Belo-sur-Mer | CAMS 142636 | 2275 | 35 | 2231 | 2339 | 2137 | Crowley [1] |
| **ATS24Hippo9** | AMNH uncatalogued | level 80+cm, layer 3 | Molar | Belo-sur-Mer | | | | | | | |
| **ATS24Hippo10** | AMNH uncatalogued | SQ 0.5-1.5, M-0, 0-16cm | | Belo-sur-Mer | | | | | | | |
| **ATS24Hippo11** | AMNH uncatalogued | SQ 2-Oh layer 2 | Vertebra | Belo-sur-Mer | | | | | | | |
| **ATS24Hippo12** | AMNH uncatalogued | A subzero SE 180-200cm | Mandible | Taolambiby | CAMS 142896 | 2905 | 40 | 2993 | 3153 | 2866 | Crowley [1] |
| **ATS24Hippo13** | AMNH uncatalogued | B subzero SE 160 cm | mandible | Taolambiby | | | | | | | |
| **ATS24Hippo14** | AMNH uncatalogued | SQB1 | Phalanx | Taolambiby | CAMS 150585 | 2620 | 110 | 2636 | 2924 | 2351 | Godfrey and Crowley [2] |
| **ATS24Hippo15†** | AMNH uncatalogued | | Mandible | Taolambiby | | | | | | | |
| **ATS24Hippo16** | NHMW C 903 | NHMW C 903 | Rib | Antsirabe | | | | | | | |
| **ATS24Hippo17** | NHMW F 906 | NHMW F 906 | Rib | Antsirabe | | | | | | | |
| **ATS24Hippo18** | NHMW D 904 | NHMW D 904 | Rib | Antsirabe | | | | | | | |
| **ATS24Hippo19** | NHMW B 902 | NHMW B 902 | Rib | Antsirabe | | | | | | | |
| **ATS24Hippo20** | NHMW A1 | NHMW A1 | Rib | Antsirabe | CAMS 142540 | 1260 | 25 | 1123 | 1259 | 1062 | Crowley [1] |
| **ATS24Hippo21** | AMNH uncatalogued | | Tooth | Anjohibe | | | | | | | |
| **ATS24Hippo22** | AMNH uncatalogued | ANJ-M-2 0-14cm | Miscellaneous teeth | Anjohibe | | | | | | | |
| **ATS24Hippo23** | AMNH uncatalogued | TW7 | Juvenile mandible | Anjohibe | β-64961/CAMS-8648 | 5300 | 60 | 6058 | 6262 | 5912 | Burney, et al. [3] |
| **ATS24Hippo24** | AMNH uncatalogued | TW7 | Neonate mandible | Anjohibe | | | | | | | |
| **ATS24Hippo25** | AMNH uncatalogued | M-1 0-9 cm | Misc bones | Anjohibe | | | | | | | |
| **ATS24Hippo26** | AMNH uncatalogued | Hippo site SQ3 | Premolar 4 from sample A1 | Anjohibe | | | | | | | |
| **ATS24Hippo27** | AMNH uncatalogued | Hippo site SQ4 | Molar 1 | Anjohibe | | | | | | | |
| **ATS24Hippo28** | UABEC 1534 | Site-1 Hole B | Femur | Antsirafaly | | | | | | | |
| **ATS24Hippo29*** | ? | DB-03-17 | Tusk #1 | No provenance | AA-45965 T-160-46A | 99 | 36 | 81 | 254 | 0 | Burney, et al. [4] |
| **ATS24Hippo30** | ? | DB-03-18 | Tooth | No provenance | CAMS 194195 | 195 | 30 | 188 | 301 | 0 | This paper |
| **ATS24Hippo32** | AMNH uncatalogued | | Radius ulna | Belo-sur-Mer | | | | | | | |
| **ATS24Hippo33** | AMNH uncatalogued | SQ 4 70-90 cm | Metatarsal | Belo-sur-Mer | β-830889 | 1930 | 30 | 1828 | 1917 | 1735 | This paper |
| **ATS24Hippo34** | AMNH uncatalogued | SQ 3-5 100+ cm | Vertebra | Belo-sur-Mer | | | | | | | |
| **ATS24Hippo35** | AMNH uncatalogued | SQ 3-5 100+ cm | Mandible | Belo-sur-Mer | | | | | | | |
| **ATS24Hippo36** | AMNH uncatalogued | SQ 3-5 70-100 cm | Partial incisor | Belo-sur-Mer | | | | | | | |
| **ATS24Hippo37** | AMNH uncatalogued | SQ. 5 Level 4 50-70 cm | Incisor | Belo-sur-Mer | | | | | | | |

| | | | | | | | | | | |
|---|---|---|---|---|---|---|---|---|---|---|
| **ATS24Hippo38** | AMNH uncatalogued | Hippo site SQ 3 | Isolated teeth/Sampled molar | Anjohibe | | | | | | |
| **ATS24Hippo39** | AMNH uncatalogued | Hippo site SQ 2 | Femur (left) | Anjohibe | | | | | | |
| **ATS24Hippo40** | AMNH uncatalogued | SQ5-6 90-110cm | Vertebra | Belo-sur-Mer | | | | | | |
| **ATS24Hippo41** | AMNH uncatalogued | SQ 5 3 35-50cm | Bone | Belo-sur-Mer | | | | | | |
| **ATS24Hippo42** | AMNH uncatalogued | no label | Molar | Belo-sur-Mer | | | | | | |
| **ATS24Hippo43** | AMNH uncatalogued | | Demimandible | Taolambiby | | | | | | |
| **ATS24Hippo44** | AMNH uncatalogued | | Mandible fragment with two teeth | Taolambiby | | | | | | |
| **ATS24Hippo45** | AMNH uncatalogued | | Mandible fragment with two teeth | Taolambiby | | | | | | |
| **ATS24Hippo46** | AMNH uncatalogued | | Tibia | Taolambiby | CAMS 143060 | 2300 | 50 | 2230 | 2352 | 2135 | Crowley [1] |
| **ATS24Hippo47** | AMNH uncatalogued | | Humerus | Taolambiby | | | | | | |
| **ATS24Hippo48** | AMNH uncatalogued | SQ 9 Layer 2 48cmbs | Molar | Belo-sur-Mer | | | | | | |
| **ATS24Hippo49** | AMNH FM 10668 | AMNH FM 10668 | Rib | Ampoza | | | | | | |
| **ATS24Hippo50** | AMNH FM 30001 | AMNH FM 30001 | Mandible | Ampoza | | | | | | |
| **ATS24Hippo51** | AMNH FM 30002 | AMNH FM 30002 | Mandible | Ampoza | | | | | | |
| **ATS24Hippo52** | AMNH FM 30003 | AMNH FM 30003 | Mandible | Ampoza | CAMS 194196 | 2535 | 40 | 2578 | 2736 | 2370 | this paper |
| **ATS24Hippo53** | AMNH FM 30004 | AMNH FM 30004 | Skull | Ampoza | β-830890 | 2340 | 30 | 2227 | 2357 | 2149 | this paper |
| **ATS24Hippo54** | AMNH FM 30006 | AMNH FM 30006 | Skull | Ampoza | | | | | | |
| **ATS24Hippo63** | Musée de l'Académie Malgache, accession unknown | DB-03-20 | | Mananjary | NZA-18525 R - 28331/10a | 2327 | 40 | 2227 | 2358 | 2142 | Burney, et al. [4] |
| **ATS24Hippo65** | AMNH uncatalogued | | Proximal scapula | Taolambiby | | | | | | |
| **ATS24Hippo66** | AMNH uncatalogued | Hippo site SQ 3 | Isolated teeth/ Sampled tusk | Anjohibe | | | | | | |
| **ATS24Misc2** | AMNH uncatalogued | ANJ-M-1 0-16cm | Miscellaneous teeth | Anjohibe | | | | | | |
| **ATS24Misc7** | AMNH uncatalogued | SQ 5-3? Bag 9 | Teeth | Belo-sur-Mer | | | | | | |

†Labelled as a Taolambiby mandible collected in the 1980s; no field or excavation number is associated with the specimen.

*Association of DB-03-17 with the published DB-HA-1 radiocarbon determination is based on specimen/sample labels and associated material; Burney et al. (2004) reports the dated specimen as DB-HA-1.

**Table S2:** Mitochondrial (mt) reference sequences used for hybridization-capture bait design. Reference accessions are provided for publicly available mt sequences. Unpublished in-house references were used solely for bait design and were not included in downstream phylogenetic analyses.

| *Species* | *NCBI Reference* |
|---|---|
| *Aepyornis maximus* | OP413809 |
| *Aldabrachelys abrupta* | OW244296 |
| *Aldabrachelys grandidieri* | OW244298 |
| *Aldabrachelys gigantea* | NC028438 |
| *Astrochelys radiata* | KJ489403 |
| *Astrochelys sp* | OW244302 |
| *Astrochelys yniphora* | JX317746 |
| *Brookesia decaryi* | AB474914 |
| *Calumma parsonii* | AB474915 |
| *Chalarodon madagascariensis* | AB266748 |
| *Crocodylus niloticus* | NC008142 |
| *Cryptoprocta ferox* | NC053961 |
| *Daubentonia madagascariensis* | AB371085 |
| *Echinops telfairi* | AB099484 |
| *Eulemur fulvus* | AB371086 |
| *Fossa fossana* | NC053959 |
| *Furcifer oustaleti* | AB185326 |
| *Furcifer petteri* | Unpublished in-house reference |
| *Choeropsis liberiensis* | NC020697 |
| *Hippopotamus amphibius* | NC000889 |
| *Lepilemur ahmansoni* | NC034733 |
| *Megaladapis edwardsi* | NC026088 |
| *Microcebus murinus* | NC028718 |
| *Microgale fotsifotsy* | BK059354 |
| *Mullerornis modestus* | OP413793 |
| *Oplurus grandidieri* | AB218720 |
| *Orycteropus afer* | NC002078 |
| *Pachylemur jullyi* | KJ944219 |
| *Palaeopropithecus ingens* | NC026090 |
| *Paroedura picta* | NC028326 |
| *Propithecus edwardsi* | KJ944243 |
| *Pyxis arachnoides* | LR697080 |
| *Pyxis planicauda* | LR697081 |
| *Uroplatus ebenaui* | AB738950 |
| *Voay robustus* | Reference assembled from public sequence data (SRA: SRR13176432) |

**Table S3: Comparative mitochondrial sequence data included in phylogenetic analyses.** Publicly available complete mitogenomes and raw-read datasets used to place the newly generated Malagasy hippopotamus sequences in a broader hippopotamid context. Sequence/BioSample and raw-read accessions, data type and original source are provided. Mitogenomes reconstructed from public raw-read datasets were generated in this study.

| Taxon | Specimen / sample | Sequence or BioSample accession | Raw-read accession(s) | Data type | Reference | Notes |
|---|---|---|---|---|---|---|
| ***Hippopotamus amphibius*** | — | NC_000889 / AJ010957 | — | Complete mitogenome | [5] | — |
| ***Hippopotamus amphibius*** | — | AP003425 | — | Complete mitogenome | — | — |
| ***Choeropsis liberiensis*** | — | NC_020697 / JN632625 | — | Complete mitogenome | [6] | — |
| ***Choeropsis liberiensis*** | — | SAMEA115628506 | ERR13148090 | Raw sequence data | — | Mitogenome reconstructed in this study |
| **European *Hippopotamus amphibius*** | NK37 | — | SRR28828336 | Raw sequence data | [7] | Mitogenome reconstructed in this study |
| **European *Hippopotamus amphibius*** | Meng19 | — | SRR28828338 | Raw sequence data | [7] | Mitogenome reconstructed in this study |
| **European *Hippopotamus amphibius*** | HgimEpp | — | SRR28828340 | Raw sequence data | [7] | Mitogenome reconstructed in this study |
| ***Hippopotamus minor*** | — | — | SRR16494613, SRR16494614, SRR14718536, SRR14718537 | Raw sequence data | [8] | Mitogenome reconstructed in this study |
| ***Hippopotamus amphibius*** | — | SAMN04009372 | SRR2183469 | RNA-seq | — | Mitogenome reconstructed in this study |
| ***Hippopotamus amphibius*** | — | SAMN07206988 | SRR5663650 | Raw sequence data | [9] | Mitogenome reconstructed in this study |
| ***Hippopotamus amphibius*** | — | SAMN10485219 | SRR8270566 | RNA-seq | — | Mitogenome reconstructed in this study |
| ***Hippopotamus amphibius*** | — | SAMN12873324 | SRR10251462 | Target capture | [10] | Mitogenome reconstructed in this study |
| ***Hippopotamus amphibius*** | — | SAMN07678089 | SRR14572852 | Raw sequence data | — | Mitogenome reconstructed in this study |
| ***Hippopotamus amphibius*** | — | SAMN07187852 | SRR5642367 | RNAseq | [11] | Mitogenome reconstructed in this study |
| ***Balaenoptera musculus*** | — | NC_001601 | — | Complete mitogenome | [12] | — |

**Table S4: Partitioning schemes and substitution models selected for mitochondrial phylogenetic analyses.** Best-fitting partition schemes and substitution models selected under the Bayesian information criterion for the full and reduced mitogenome datasets using either *Choeropsis liberiensis* or *Balaenoptera musculus* as outgroup.

| | | | |
|---|---|---|---|
| | | **_Choeropsis liberiensis_ outgroup full dataset** | |
| Partition | Best Model | #sites | Positions |
| 1 | HKY+I | 6160 | tRNAs, 16S, 12S, Intergenic regions, first codon position of ND1, ND2, ND4, ND5, ATP6, ATP8, and second codon position of ATP8 |
| 2 | TRN+I | 3560 | Second codon position of CO2, CYTB, ND5, CO3, CO1, ND1, ND2, ND4, ND3, ATP6, ND4L |
| 3 | TRN+G | 3611 | Third codon position of ND5, CYTB, ND3, ND1, ATP6, ND2, ND4, CO3, CO1, ND4L, CO2, ATP8 |
| 4 | K80+I | 1600 | First codon position of ND3, CO2, CYTB, CO1, CO3, ND4L |
| 5 | HKY+I | 343 | First and second codon position of ND6 |
| 6 | TRN | 171 | Third codon position of ND6 |
| 7 | TRN+I+G | 963 | Control Region |
| | | **_Choeropsis liberiensis_ outgroup reduced dataset** | |
| Partition | Best Model | #sites | Positions |
| 1 | HKY+I | 6980 | tRNAs, 16S, 12S, Intergenic regions, first codon position of ND5, ATP6, ND1, ND2, ND4, ATP8, , ND4L, ND3, CO2, CYTB, second codon position of ATP8 |
| 2 | HKY+I | 3731 | Second codon position of ATP6, ND4L, ND2, ND3, ND1, ND6, ND4, CO3, CYTB, ND5, CO1, CO2 |
| 3 | TRN+I | 3611 | Third codon position of ND3, ND5, CYTB, ND1, ND2, ATP6, ND4, CO3, CO1, ND4L, CO2, ATP8 |
| 4 | K80+I | 780 | First codon position of CO1, CO3 |
| 5 | TRN+G | 343 | First and third codon position of ND6 |
| 6 | TRN+I+G | 963 | Control Region |
| | | **_Balaenoptera musculus_ outgroup full dataset** | |
| Partition | Best Model | #sites | Positions |
| 1 | TRN+G | 4177 | tRNAs, 16S, 12S, first codon position of ND4L |
| 2 | K80+I | 1824 | First codon position of ND3, ND1, CO1, CO2, CO3, CYTB |
| 3 | TRN+I | 3733 | Second codon position of ATP6, ND2, ND3, ND5, ND6, CYTB, ND4L, ND4, ND1, CO1, CO3, CO2 |
| 4 | TRN+I+G | 3612 | Third codon position of ND3, ND2, ND1, CYTB, ND5, CO1, ND4L, ATP8, CO2, ND4, CO3, ATP6 |
| 5 | HKY+I | 1803 | Intergenic regions, first codon position of ATP8, ND4, ND2, ND5, ATP6, second codon position of ATP8 |
| 8 | TRN+G | 172 | First codon position of ND6 |
| 9 | TRN | 171 | Third codon position of ND6 |
| | | **_Balaenoptera musculus_ outgroup reduced dataset** | |
| Partition | Best Model | #sites | Positions |
| 1 | TRN+G | 4177 | tRNAs, 16S, 12S, first codon position of ND4L |
| 2 | HKY+I | 1043 | First codon position of ND3, ND1, CO2, CYTB |
| 3 | HKY+I | 2724 | Second codon position of ATP6, ND2, ND3, ND6, ND5, CYTB, ND4L, ND4, ND1 |
| 4 | TRN+I+G | 3612 | Third codon position of ND3, ND2, ND1, CYTB, ND5, CO1, ND4L, ATP8, CO2, ND4, CO3, ATP6 |
| 5 | HKY+I | 1803 | Intergenic regions, first codon position of ATP8, ND4, ND2, ND5, ATP6, second codon position of ATP8 |
| 6 | TRNEF+I | 781 | First codon position of CO1, CO3 |
| 7 | TRN+I | 1009 | Second codon position of CO1, CO3, CO2 |
| 8 | TRN+G | 172 | First codon position of ND6 |
| 9 | TRN | 171 | Third codon position of ND6 |

**Table S5: Tip-age information and priors used in Bayesian divergence-time analyses.** Radiocarbon-dated specimens were assigned fixed tip ages corresponding to the median calibrated radiocarbon age. Undated specimens were assigned uniform age priors based on locality-specific chronological information; prior bounds and their supporting sources are provided.

| Locality | Uniform tip-date prior (cal BP) | Chronological basis | References |
|---|---|---|---|
| **Belo-sur-Mer** | 900–3,000 | Late-Holocene chronology for Belo-sur-Mer/Ankilibehandry; published hippopotamus material includes dates around 2.1–2.3 ka, with bounds broadened to encompass the local subfossil chronology | Godfrey and Crowley [2], Burney, et al. [3], Burney, et al. [4], Crowley, et al. [13] |
| **Taolambiby** | 1,000–3,500 | Published directly dated hippopotamus specimens span approximately 1.27–3.13 ka cal BP; bounds rounded outward | Godfrey and Crowley [2], Crowley, et al. [13] |
| **Anjohibe** | 2,400–7,500 | Published hippopotamus dates span approximately 2.49–7.31 ka cal BP; bounds rounded outward to encompass the known site chronology | Godfrey and Crowley [2], Burney, et al. [4], Crowley, et al. [13] |
| **Antsirafaly** | 900–2,500 | Directly dated Antsirafaly fauna spans roughly 0.9–1.8 ka cal BP, including *H. lemerlei* at ~1.12 ka; a broader conservative upper bound was used | Godfrey, et al. [14] |
| **Ampoza** | 1,000–3,500 | Published Ampoza hippopotamus dates fall within the late Holocene, including specimens around ~1.2–2.8 ka; bounds broadened and rounded to encompass the known local chronology | Godfrey and Crowley [2], Burney, et al. [4], Crowley, et al. [13], Hansford, et al. [15] |

**Table S6: Stable isotope data used in ecological comparisons.** Stable carbon ($\delta^{13}C$) and nitrogen ($\delta^{15}N$) isotope values for Malagasy hippopotamus specimens from localities represented in the genetic dataset. Specimen locality, mitochondrial lineage where available, isotope values and original data source are provided. Isotope values newly generated in this study are identified as "This study" in the source columns.

| Locality | ATS# | Specimen No | d13C | d15N | d13C Source | d15N Source | Lineage |
|---|---|---|---|---|---|---|---|
| **Antsirabe** | ATS24Hippo20 | NHMW uncatalogued | -25.45 | 2.89 | Godfrey and Crowley [2] | Godfrey and Crowley [2] | Antsirabe/Anjohibe lineage |
| **Antsirabe** | – | NHMW uncatalogued | -25.01 | 2.96 | Godfrey and Crowley [2] | Godfrey and Crowley [2] | – |
| **Antsirabe** | – | NHMW uncatalogued | -24.9 | 1.89 | Godfrey and Crowley [2] | Godfrey and Crowley [2] | – |
| **Antsirabe** | – | NHMUK PV M7093 | -25.4 | 3.8 | Hansford and Turvey [16] | Hansford and Turvey [16] | – |
| **Antsirabe** | – | NHMUK PV M5142 | -21.9 | 4.1 | Hansford and Turvey [16] | Hansford and Turvey [16] | – |
| **Antsirabe** | – | NHMUK PV M5141 | -18.9 | 3 | Hansford and Turvey [16] | Hansford and Turvey [16] | – |
| **Antsirabe** | – | NHMW uncatalogued | -25.94 | 6.49 | Godfrey and Crowley [2] | Godfrey and Crowley [2] | – |
| **Antsirabe** | – | NHMW uncatalogued | -23.67 | 1.34 | Godfrey and Crowley [2] | Godfrey and Crowley [2] | – |
| **Anjohibe** | – | UA uncatalogued | -22.08 | 11.47 | Crowley and Samonds [17] | Crowley, et al. [18] | – |
| **Anjohibe** | – | UA uncatalogued | -22.51 | 5.45 | Crowley and Samonds [17] | Crowley, et al. [18] | – |
| **Anjohibe** | – | UA 3591 | -24.29 | 8.06 | Crowley and Samonds [17] | Crowley, et al. [13] | – |
| **Anjohibe** | – | UA 9570 | -24.13 | 6.54 | Crowley and Samonds [17] | Crowley, et al. [13] | – |
| **Anjohibe** | – | UA uncatalogued | -23.05 | 5.59 | Crowley and Samonds [17] | Crowley, et al. [13] | – |
| **Anjohibe** | – | UA 4917 | -22.97 | 7.47 | Crowley and Samonds [17] | Crowley, et al. [13] | – |
| **Anjohibe** | – | UA 3560 | -21.59 | 7.33 | Crowley and Samonds [17] | Crowley, et al. [13] | – |
| **Anjohibe** | – | UA 3558 | -21.14 | 7.24 | Crowley and Samonds [17] | Crowley, et al. [13] | – |
| **Anjohibe** | – | UA uncatalogued | -20.96 | 6.41 | Crowley and Samonds [17] | Crowley, et al. [13] | – |
| **Antsirafaly** | – | UABEC 1377 | -15.97 | 10.44 | Crowley, et al. [18] | Crowley, et al. [18] | – |
| **Antsirafaly** | – | UABEC 1378 | -13.15 | 10.87 | Crowley, et al. [18] | Crowley, et al. [18] | – |
| **Antsirafaly** | – | UABEC 1391 | -12.77 | 11.90 | Crowley, et al. [18] | Crowley, et al. [18] | – |
| **Antsirafaly** | – | UABEC 1455 | -16.45 | 12.03 | Crowley, et al. [18] | Crowley, et al. [18] | – |
| **Antsirafaly** | – | UABEC 1453 | -15.51 | 11.56 | Crowley, et al. [18] | Crowley, et al. [18] | – |
| **Antsirafaly** | – | UABEC 1456 | -15.30 | 11.20 | Crowley, et al. [18] | Crowley, et al. [18] | – |
| **Antsirafaly** | – | UABEC 1381 | -14.66 | 12.63 | Crowley, et al. [18] | Crowley, et al. [18] | – |
| **Antsirafaly** | – | UABEC 1472 | -14.52 | 11.58 | Crowley, et al. [18] | Crowley, et al. [18] | – |

| | | | | | | | |
|---|---|---|---|---|---|---|---|
| **Antsirafaly** | – | UABEC 1376 | -14.13 | 12.27 | Crowley, et al. [18] | Crowley, et al. [18] | – |
| **Antsirafaly** | – | UABEC 1375 | -13.68 | 12.02 | Crowley, et al. [18] | Crowley, et al. [18] | – |
| **Antsirafaly** | – | UABEC 1429 | -12.96 | 11.99 | Crowley, et al. [18] | Crowley, et al. [18] | – |
| **Antsirafaly** | – | UABEC 1379 | -13.81 | 10.37 | Crowley, et al. [18] | Crowley, et al. [18] | – |
| **Taolambiby** | – | Not reported | -19.32 | 9.8 | Anderson, et al. [19] | Hixon, et al. [20] | – |
| **Taolambiby** | – | Not reported | -17.62 | 9 | Anderson, et al. [19] | Hixon, et al. [20] | – |
| **Taolambiby** | – | Not reported | -17.12 | 8.6 | Anderson, et al. [19] | Hixon, et al. [20] | – |
| **Taolambiby** | – | UM uncatalogued | -20.28 | 13.14 | Crowley and Godfrey [21] | Crowley and Godfrey [21] | – |
| **Taolambiby** | – | UM uncatalogued | -19.99 | 8.61 | Crowley and Godfrey [21] | Crowley and Godfrey [21] | – |
| **Taolambiby** | ATS24Hippo12 | AMNH uncatalogued | -18.61 | 8.43 | Crowley and Godfrey [21] | Crowley and Godfrey [21] | No DNA |
| **Taolambiby** | – | UM uncatalogued | -17.63 | 10.71 | Crowley and Godfrey [21] | Crowley and Godfrey [21] | – |
| **Taolambiby** | – | UM uncatalogued | -17.25 | 6.75 | Crowley and Godfrey [21] | Crowley and Godfrey [21] | – |
| **Taolambiby** | – | UM uncatalogued | -17.1 | 7.66 | Crowley and Godfrey [21] | Crowley and Godfrey [21] | – |
| **Taolambiby** | ATS24Hippo46 | AMNH uncatalogued | -18.92 | 7.31 | Crowley and Godfrey [21] | Crowley and Godfrey [21] | Taolambiby associated lineage |
| **Taolambiby** | – | UM uncatalogued | -18.08 | 8.75 | Crowley and Godfrey [21] | Crowley and Godfrey [21] | – |
| **Taolambiby** | – | | -21 | 9 | Hixon, et al. [20] | Hixon, et al. [20] | – |
| **Taolambiby** | – | | -20.3 | 10.1 | Hixon, et al. [20] | Hixon, et al. [20] | – |
| **Taolambiby** | – | | -19.7 | 9.6 | Hixon, et al. [20] | Hixon, et al. [20] | – |
| **Taolambiby** | – | | -19.5 | 9.7 | Hixon, et al. [20] | Hixon, et al. [20] | – |
| **Taolambiby** | – | | -17.7 | 8.9 | Hixon, et al. [20] | Hixon, et al. [20] | – |
| **Ampoza** | – | UA uncatalogued | -21.57 | 11.82 | Burney, et al. [4] | Crowley and Godfrey [21] | – |
| **Ampoza** | – | UA uncatalogued (B3-13) | -22.4 | 8.96 | Crowley and Godfrey [21] | Crowley and Godfrey [21] | – |
| **Ampoza** | – | PMU 3972 | -22.5 | 9.6 | Hansford and Turvey [16] | Hansford and Turvey [16] | – |
| **Ampoza** | – | PMU 3973 | -22.5 | 9.3 | Hansford and Turvey [16] | Hansford and Turvey [16] | – |
| **Ampoza** | – | NHMUK PV M 82190 | -21.7 | 9.7 | Hansford and Turvey [16] | Hansford and Turvey [16] | – |
| **Ampoza** | – | MNHN MAD7918 | -21 | 7 | Hansford and Turvey [16] | Hansford and Turvey [16] | – |
| **Ampoza** | – | UA uncatalogued (B3-117-P) | -22.08 | 7.56 | Crowley and Godfrey [21] | Crowley and Godfrey [21] | – |
| **Ampoza** | – | UA10621, B3-185 | -24.3 | 9.4 | Hixon, et al. [20] | Hixon, et al. [20] | – |
| **Ampoza** | – | UA10635, E1-53 | -23.1 | 9.5 | Hixon, et al. [20] | Hixon, et al. [20] | – |
| **Ampoza** | – | PMU 32160 | -23 | 9.2 | Hixon, et al. [20] | Hixon, et al. [20] | – |
| **Ampoza** | – | UA10617, B3-5 | -22.9 | 8.6 | Hixon, et al. [20] | Hixon, et al. [20] | – |

| | | | | | | | |
|---|---|---|---|---|---|---|---|
| **Ampoza** | – | UA10623, A4-11 | -22.8 | 9.5 | Hixon, et al. [20] | Hixon, et al. [20] | – |
| **Ampoza** | – | UA uncatalogued (A4-127) | -22.6 | 10.1 | Hixon, et al. [20] | Hixon, et al. [20] | – |
| **Ampoza** | – | UA10622, A4-12 | -22.6 | 9.5 | Hixon, et al. [20] | Hixon, et al. [20] | – |
| **Ampoza** | – | UA10628, B3-6 | -22.6 | 9.4 | Hixon, et al. [20] | Hixon, et al. [20] | – |
| **Ampoza** | – | UA10627, E1-56 | -22.4 | 10.5 | Hixon, et al. [20] | Hixon, et al. [20] | – |
| **Ampoza** | – | UA10637, B3-7 | -22.3 | 9.5 | Hixon, et al. [20] | Hixon, et al. [20] | – |
| **Ampoza** | – | UA10639, B2-20 | -22.3 | 9.9 | Hixon, et al. [20] | Hixon, et al. [20] | – |
| **Ampoza** | – | | -22.2 | 10.2 | Hixon, et al. [20] | Hixon, et al. [20] | – |
| **Ampoza** | – | UA10613, E1-55 | -22.2 | 9.6 | Hixon, et al. [20] | Hixon, et al. [20] | – |
| **Ampoza** | – | UA10619, B3-223 | -22.2 | 10.3 | Hixon, et al. [20] | Hixon, et al. [20] | – |
| **Ampoza** | – | UA10625, CG-76 | -22.2 | 9.1 | Hixon, et al. [20] | Hixon, et al. [20] | – |
| **Ampoza** | – | UA10640, A4-125 | -22 | 9.4 | Hixon, et al. [20] | Hixon, et al. [20] | – |
| **Ampoza** | – | UA10615, E1-84 | -21.5 | 9.9 | Hixon, et al. [20] | Hixon, et al. [20] | – |
| **Ampoza** | – | UA uncatalogued | -22.79 | 8.98 | Crowley and Godfrey [21] | Crowley and Godfrey [21] | – |
| **Ampoza** | – | UA uncatalogued | -22.02 | 8.66 | Crowley and Godfrey [21] | Crowley and Godfrey [21] | – |
| **Ampoza** | ATS24Hippo53 | AMNH FM 30004 | -22.67 | 9.96 | This study | This study | Belo-associated lineage |
| **Belo-sur-Mer** | ATS24Hippo33 | AMNH uncatalogued | -15.45 | 7.39 | This study | This study | Ampoza-associated lineage |
| **Ampoza** | ATS24Hippo52 | AMNH FM 30003 | -22.47 | 11.62 | This study | This study | Ampoza-associated lineage |
| **Belo-sur-Mer** | ATS24Hippo4 | AMNH uncatalogued | -14.93 | 7.77 | This study | This study | Belo-associated lineage |
| **Belo-sur-Mer** | ATS24Hippo8 | AMNH uncatalogued | -19.9 | 9.3 | This study | This study | Belo-associated lineage |

**Table S7. Fossil tip-age priors used for morphology-only terminals in the total-evidence analysis.** Fossil terminals represent species- or locality-level composite morphological OTUs derived from Boisserie, et al. [22]. Where the material underlying an OTU could be associated with a particular locality or stratigraphic unit, the narrowest published age interval was used. For more broadly composite OTUs, published stratigraphic ranges were used as conservative proxies. Ages are given in millions of years before present (Ma). Molecular subfossil terminals with specimen-specific ages are not included.

| Taxon | Principal material / age basis | Tip-age prior | Reference |
|---|---|---|---|
| ***Saotherium* cf. *mingoz*** | Kossom Bougoudi, Chad | Normal (μ = 5.26 Ma, σ = 0.29 Ma) | Lebatard, et al. [23] |
| ***Saotherium mingoz*** | Kollé, Chad | Uniform (4.00–5.00 Ma) | Lebatard, et al. [23]; Boisserie, et al. [24] |
| ***Archaeopotamus harvardi*** | Lothagam, Kenya; Nawata Formation | Uniform (5.00–7.44 Ma) | Weston [25]; McDougall and Feibel [26] |
| **aff. *Hippopotamus* cf. sp. Hadar** | Kanapoi, Kenya | Uniform (4.108–4.195 Ma) | Boisserie [27] |
| **aff. *Hippopotamus afarensis*** | Hadar Formation, Ethiopia | Uniform (2.90–3.42 Ma) | Reed [28]; Campisano and Feibel [29] |
| ***Hexaprotodon bruneti*** | Bouri/upper Maka, Ethiopia | Fixed at 2.50 Ma | Boisserie and White [30] |
| **aff. *Hippopotamus protamphibius*** | Turkana Basin | Uniform (1.87–3.60 Ma) | Harris [31]; Bobe, et al. [32] |
| **aff. *Hippopotamus aethiopicus*** | Turkana Basin | Uniform (1.38–2.60 Ma) | Harris [31]; Bobe, et al. [32] |
| **aff. *Hippopotamus karumensis*** | Turkana Basin | Uniform (1.38–2.60 Ma) | Harris [31]; Bobe, et al. [32] |
| ***Hippopotamus gorgops*** | Olduvai and Turkana Basin | Uniform (0.70–2.60 Ma) | Harris [31]; Rowan, et al. [33] |
| ***Hexaprotodon sivalensis*** | Siwalik material, South Asia | Uniform (2.20–5.90 Ma) | Barry, et al. [34] |
| ***Hexaprotodon palaeindicus*** | Narmada material, India | Uniform (0.0117–0.129 Ma) | Turvey, et al. [35] |

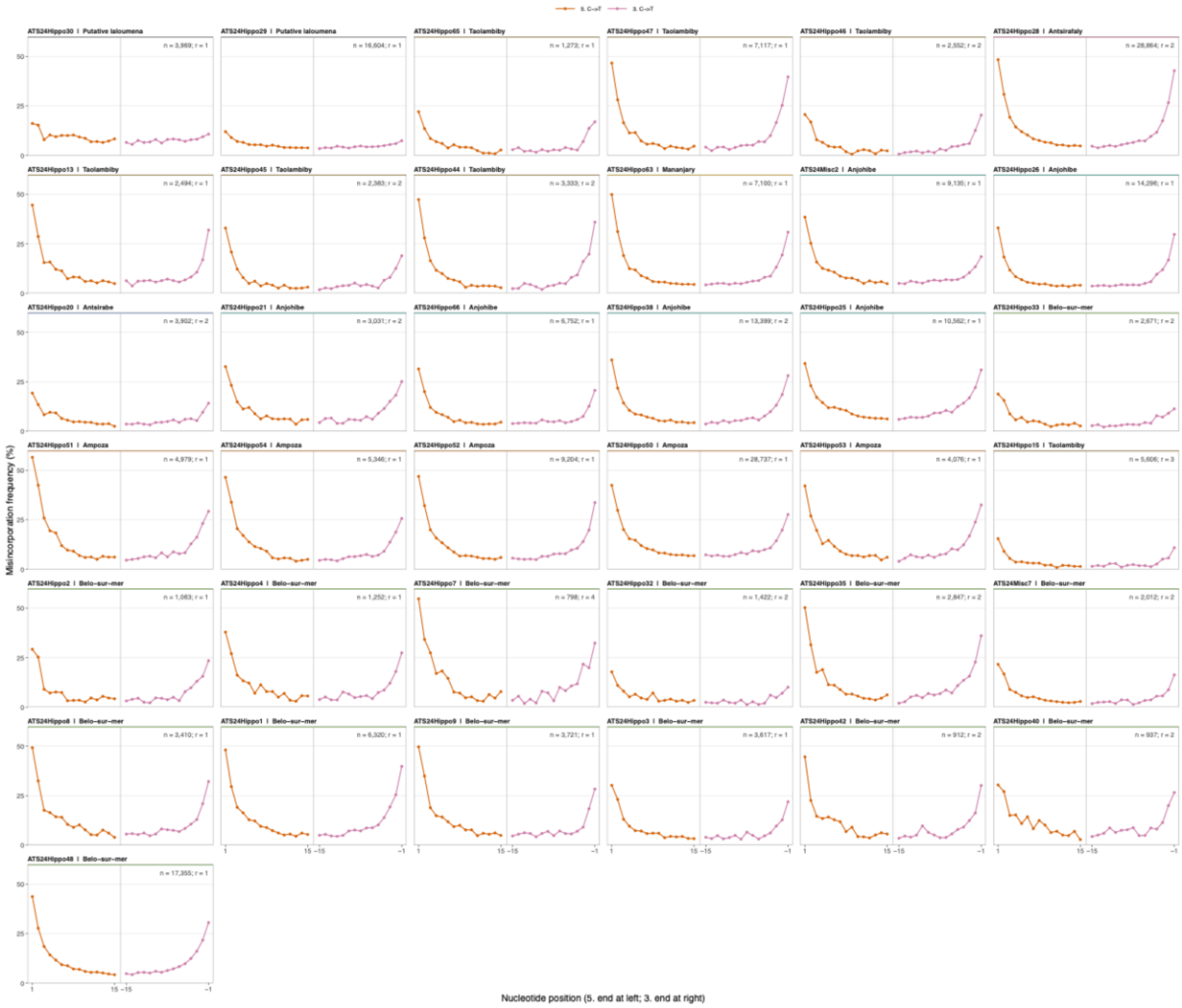


**Figure S1: Ancient DNA damage profiles for Malagasy hippopotamus specimens.** DamageProfiler estimates of nucleotide misincorporation frequencies at sequence termini for specimens included in the phylogenetic analyses. Elevated C→T substitutions toward both the 5′ and 3′ ends are characteristic of post-mortem ancient DNA damage in single-stranded libraries. Here, n denotes the number of mapped reads after deduplication, and r denotes the number of libraries per sample.

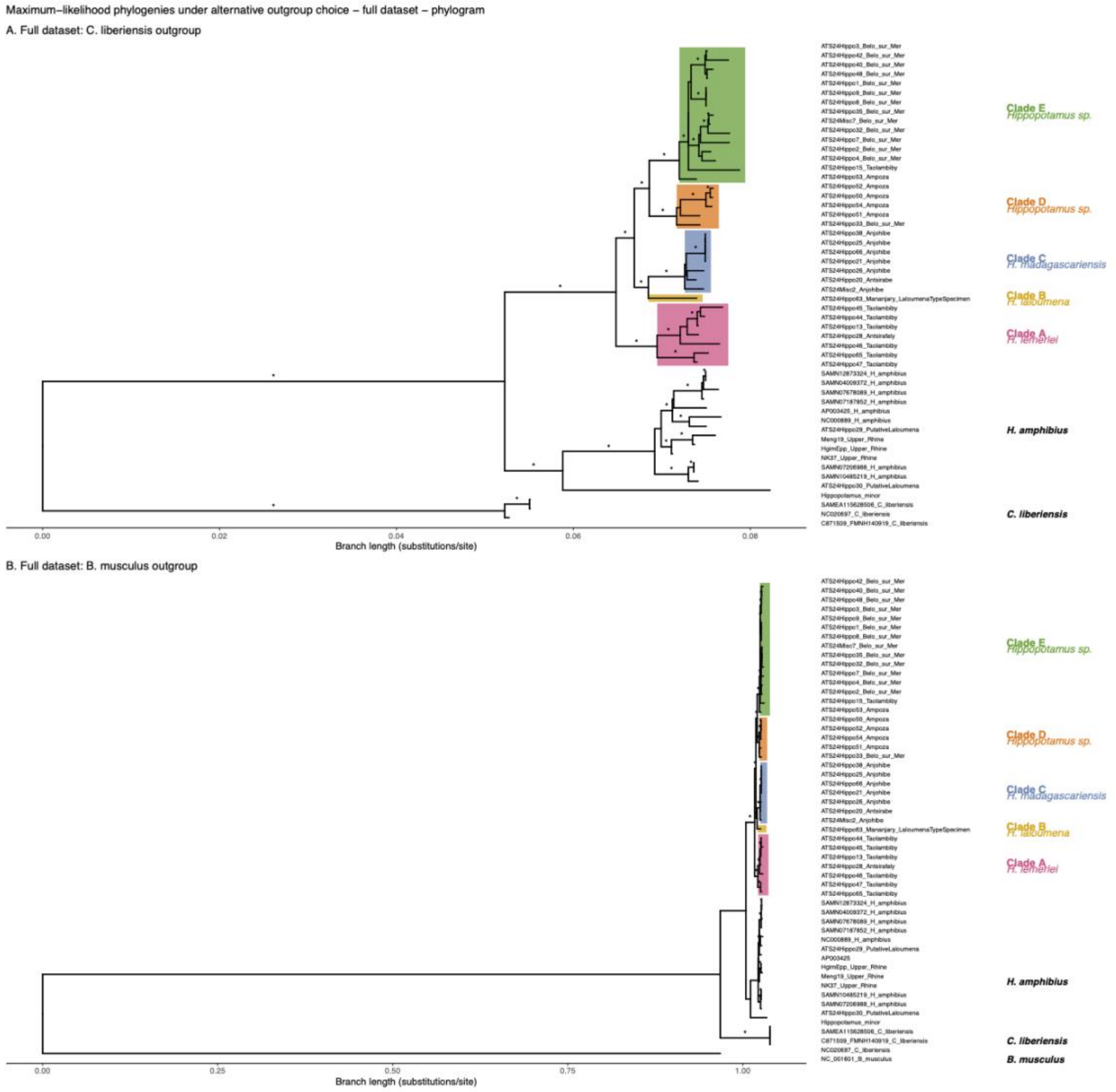


**Figure S2: Maximum-likelihood phylogenies of the full Malagasy hippopotamus mitogenome dataset under alternative outgroup treatments. (A)** Phylogeny rooted with the extant pygmy hippopotamus, *Choeropsis liberiensis*. **(B)** Phylogeny including the blue whale, *Balaenoptera musculus*, as a more distant cetacean outgroup. Tip labels identify individual specimens and comparative sequences. Bootstrap support values are shown at nodes, with asterisks indicating bootstrap support ≥90%. Both outgroup treatments recovered the same major relationships among the endemic Malagasy mt lineages.

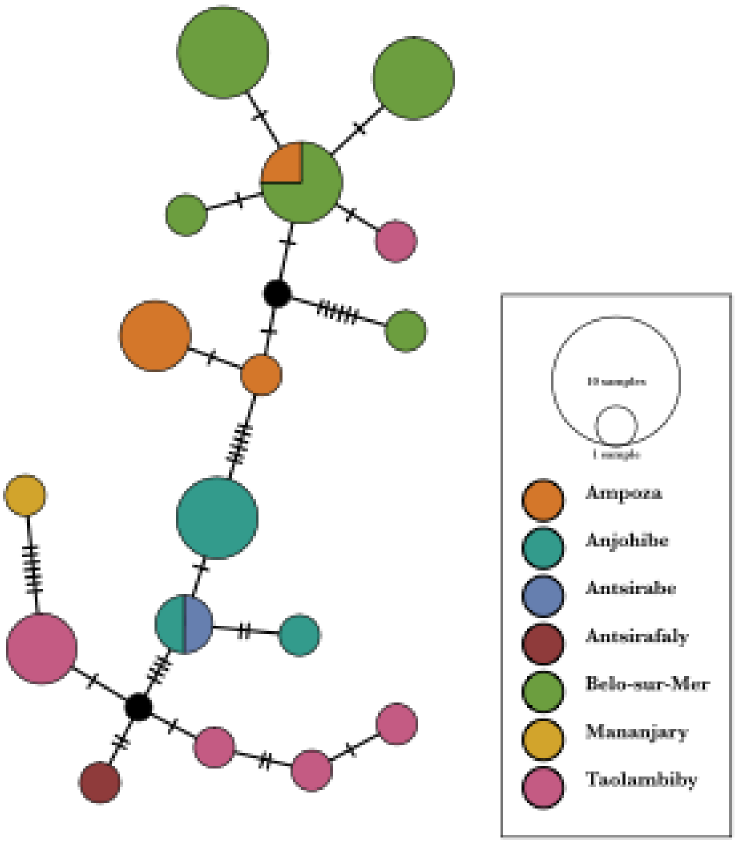


**Figure S3: Median-joining haplotype network of the broader Malagasy hippopotamus dataset.** Median-joining network constructed from confirmed endemic Malagasy specimens with >60% mitogenome coverage. Thirty-five sequences collapsed into 17 haplotypes. Circle size is proportional to haplotype frequency, colours indicate sampling locality, and hatch marks indicate mutational steps between haplotypes. The two specimens grouping with modern *Hippopotamus amphibius* were excluded.

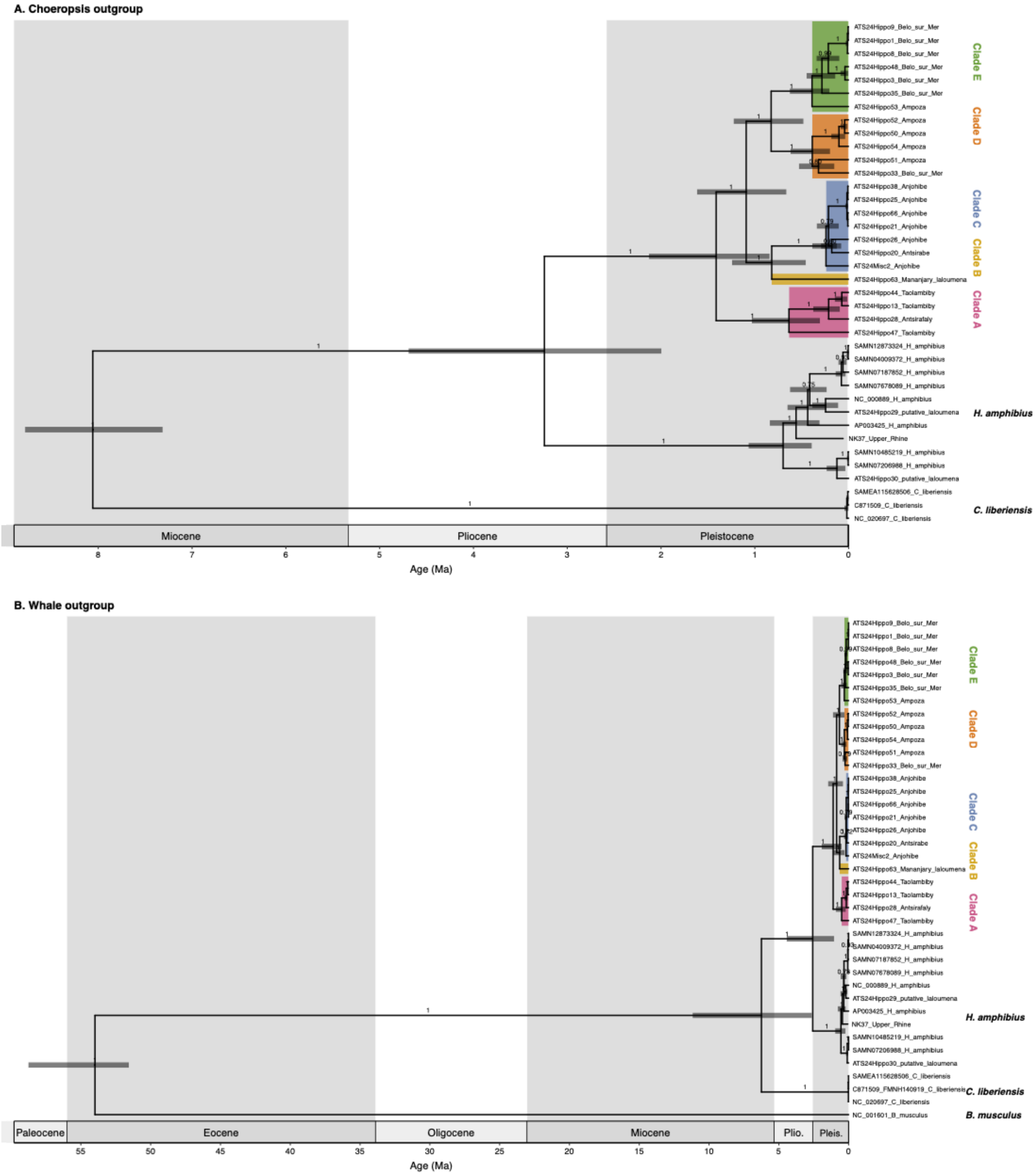


**Figure S4: Bayesian tip-dated phylogenies of the reduced high-coverage Malagasy hippopotamus mitogenome dataset under alternative outgroup and calibration schemes.** **(A)** Maximum clade credibility tree rooted with *Choeropsis liberiensis* and calibrated using the *Choeropsis–Hippopotamus* divergence. **(B)** Maximum clade credibility tree including *Balaenoptera musculus* and calibrated using the whale–hippopotamid divergence. Tip labels identify individual specimens and comparative sequences, and node bars indicate 95% highest posterior density intervals for divergence-time estimates. The two calibration schemes recovered broadly similar divergence-time estimates and the same major relationships among Malagasy lineages.

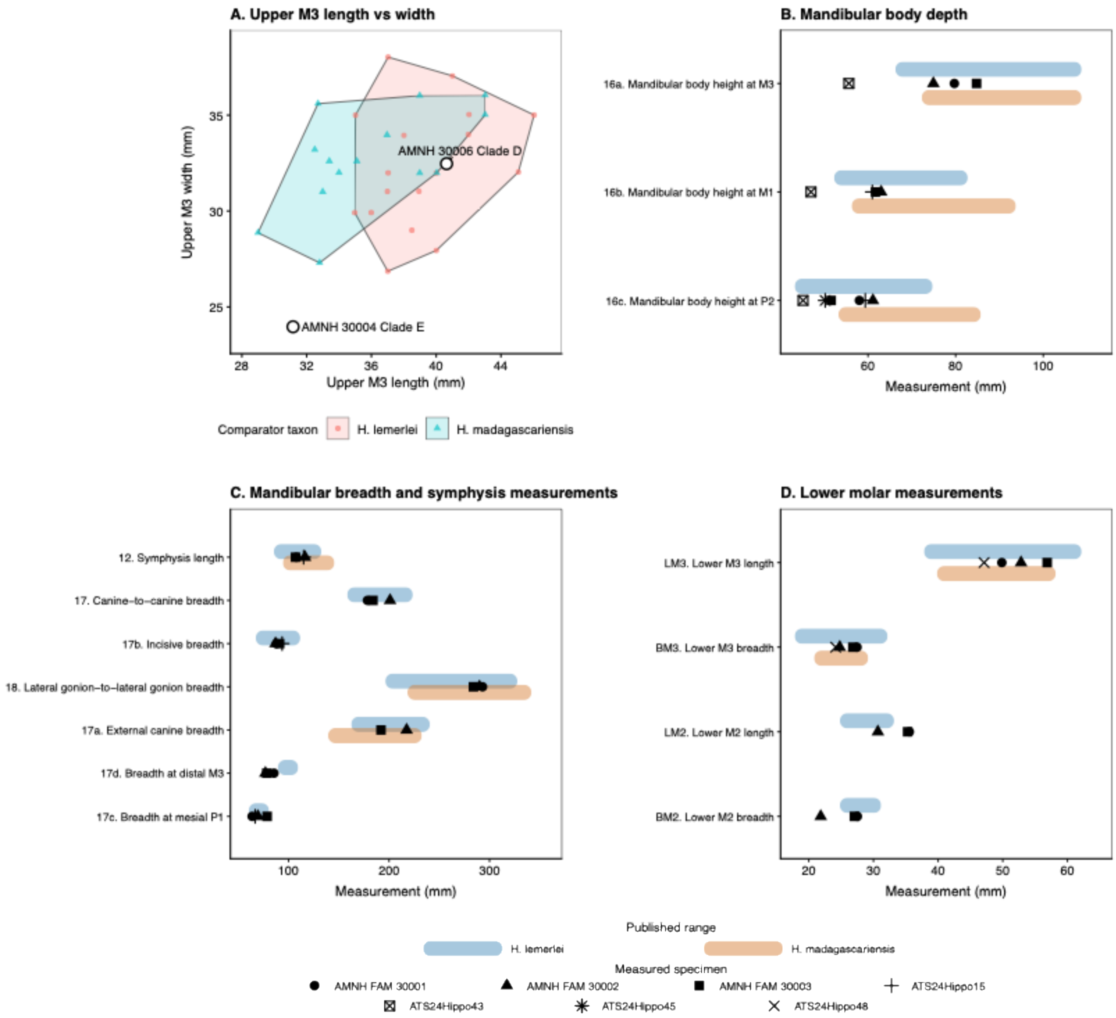


**Figure S5: Mandibular and dental comparisons of Malagasy hippopotamuses.** Measurements from genetically sampled Malagasy hippopotamus specimens compared with published measurement ranges for *H. lemerlei* and *H. madagascariensis*. Comparisons include upper third molar length and width (A), mandibular body depth (B), mandibular breadth and symphysis measurements (C), and lower molar measurements (D). Considerable overlap between the published species ranges limits the taxonomic resolution of these characters.

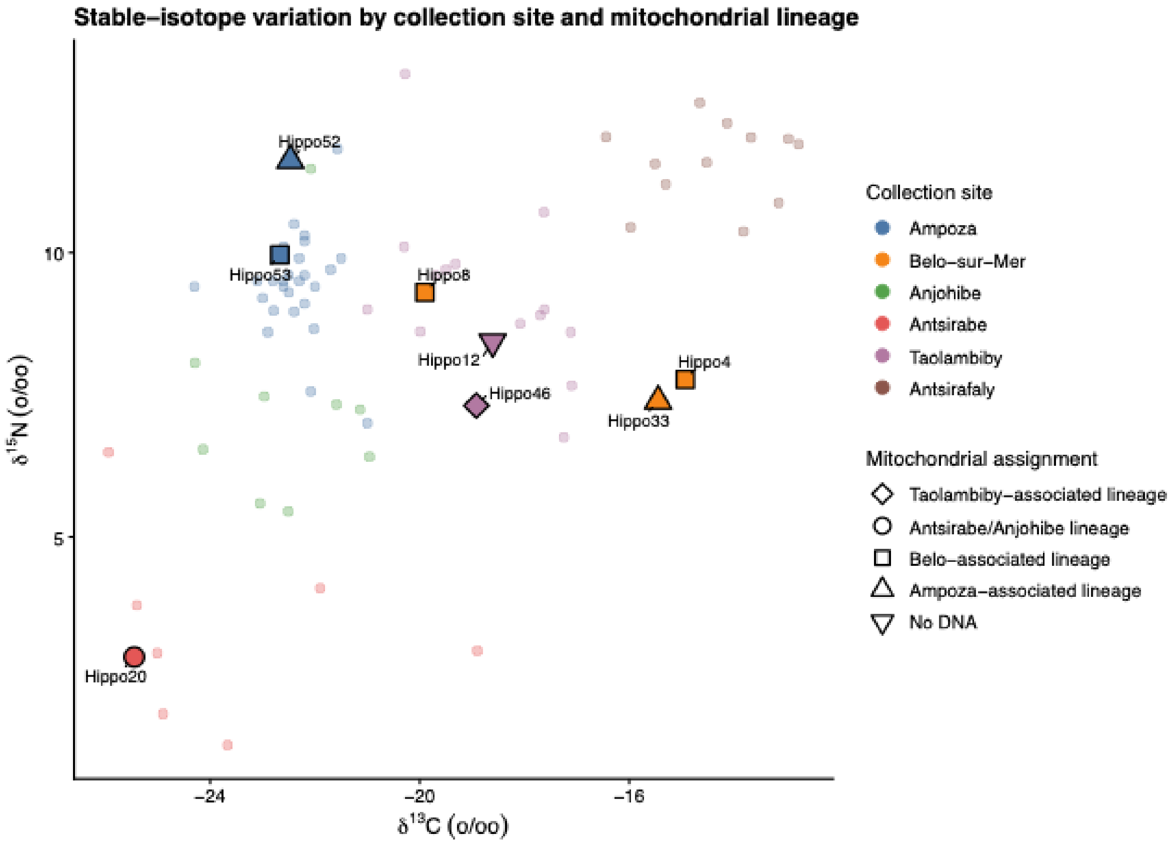


**Figure S6: Stable carbon and nitrogen isotope variation in Malagasy hippopotamuses.** Stable carbon ($\delta^{13}C$) and nitrogen ($\delta^{15}N$) isotope values for Malagasy hippopotamuses from localities represented in the genetic dataset. The dataset combines previously published isotope values with isotope values newly generated in this study. Genetically sampled specimens are identified by their mt lineage where available. Isotopic variation shows stronger geographic structuring by locality than correspondence with the major mtDNA lineages.

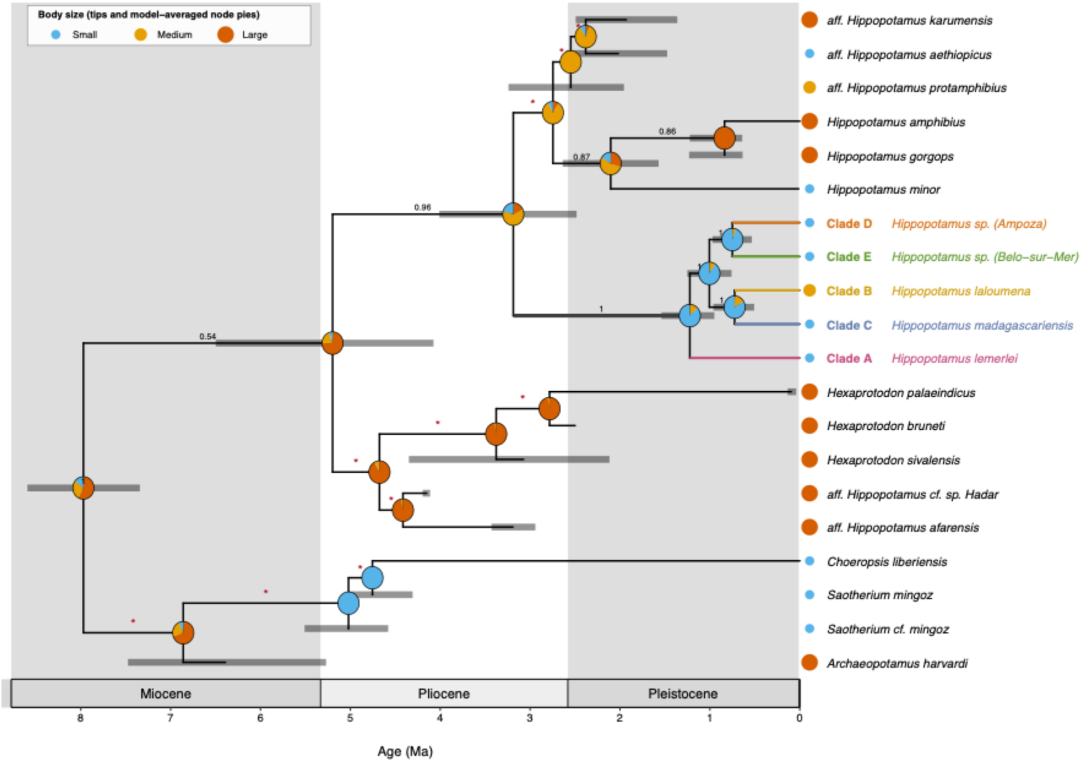


**Figure S7: Total-evidence phylogeny and ancestral reconstruction of hippopotamid body size.** Bayesian total-evidence phylogeny combining mt sequence data with 37 morphological characters for extant and extinct hippopotamids. Malagasy Clades A–E are shown within the broader hippopotamid phylogeny. Ancestral body-size states were reconstructed as small, medium or large using AICc-weighted averaging across five transition models. Pie charts show the relative support for alternative body-size states at reconstructed ancestral nodes. Numbers indicate clade posterior probabilities, with red asterisks marking nodes with posterior probabilities below 0.5. Grey horizontal bars indicate 95% highest posterior density intervals for estimated ages. The Malagasy crown ancestor is reconstructed as small-bodied, whereas the immediately preceding ancestral node favours a medium-to-large body size. The nested position of *H. laloumena* is consistent with a secondary increase in body size within the Malagasy radiation.

**Supplementary References:**


1. Crowley, B.E. A refined chronology of prehistoric Madagascar and the demise of the megafauna. *Quat. Sci. Rev.* **29**, 2591–2603 (2010).
2. Godfrey, L.R. & Crowley, B.E. Madagascar's ephemeral palaeo-grazer guild: who ate the ancient C4 grasses? *Proc. R. Soc. B* **283**(2016).
3. Burney, D.A. *et al.* Environmental change, extinction and human activity: evidence from caves in NW Madagascar. *J. Biogeogr.* **24**, 755–767 (1997).
4. Burney, D.A. *et al.* A chronology for late prehistoric Madagascar. *J. Hum. Evol.* **47**, 25–63 (2004).
5. Ursing, B.M. & Arnason, U. Analyses of mitochondrial genomes strongly support a hippopotamus-whale clade. *Proc. R. Soc. B* **265**, 2251–5 (1998).
6. Hassanin, A. *et al.* Pattern and timing of diversification of Cetartiodactyla (Mammalia, Laurasiatheria), as revealed by a comprehensive analysis of mitochondrial genomes. *C. R. Biol.* **335**, 32–50 (2012).
7. Arnold, P. *et al.* Ancient DNA and dating evidence for the dispersal of hippos into central Europe during the last glacial. *Curr. Biol.* **35**, 5363–5371 (2025).
8. Psonis, N. *et al.* Mitochondrial sequences of the extinct Cypriot pygmy hippopotamus confirm its phylogenetic placement. *Zool. J. Linn. Soc.* **196**, 979–989 (2022).
9. Arnason, U., Lammers, F., Kumar, V., Nilsson, M.A. & Janke, A. Whole-genome sequencing of the blue whale and other rorquals finds signatures for introgressive gene flow. *Sci. Adv.* **4**, eaap9873 (2018).
10. McGowen, M.R. *et al.* Phylogenomic Resolution of the Cetacean Tree of Life Using Target Sequence Capture. *Syst. Biol.* **69**, 479–501 (2020).
11. Figuet, E., Ballenghien, M., Lartillot, N. & Galtier, N. Reconstruction of body mass evolution in the Cetartiodactyla and mammals using phylogenomic data. *Peer Community J.* **1**(2021).
12. Arnason, U. & Gullberg, A. Comparison between the complete mtDNA sequences of the blue and the fin whale, two species that can hybridize in nature. *J Mol Evol* **37**, 312–22 (1993).
13. Crowley, B.E. *et al.* Island-wide aridity did not trigger recent megafaunal extinctions in Madagascar. *Ecography* **40**, 901–912 (2017).
14. Godfrey, L.R. *et al.* Teasing apart impacts of human activity and regional drought on Madagascar's large vertebrate fauna: insights from new excavations at Tsimanampesotse and Antsirafaly. *Front. Ecol. Evol.* **9**, 742203 (2021).
15. Hansford, J.P., Lister, A.M., Weston, E.M. & Turvey, S.T. Simultaneous extinction of Madagascar's megaherbivores correlates with late Holocene human-caused landscape transformation. *Quat. Sci. Rev.* **263**, 106996 (2021).
16. Hansford, J.P. & Turvey, S.T. Dietary isotopes of Madagascar's extinct megafauna reveal holocene browsing and grazing guilds. *Biol. Lett.* **18**, 20220094 (2022).
17. Crowley, B.E. & Samonds, K.E. Stable carbon isotope values confirm a recent increase in grasslands in northwestern Madagascar. *Holocene* **23**, 1066–1073 (2013).
18. Crowley, B.E., Godfrey, L.R. & Samonds, K.E. What can hippopotamus isotopes tell us about past distributions of C4 grassy biomes on Madagascar? *Plants People Planet* **5**, 997–1010 (2023).
19. Anderson, A. *et al.* New evidence of megafaunal bone damage indicates late colonization of Madagascar. *PLOS ONE* **13**, e0204368 (2018).
20. Hixon, S.W. *et al.* Late Holocene spread of pastoralism coincides with endemic megafaunal extinction on Madagascar. *Proc. R. Soc. B* **288**, 20211204 (2021).
21. Crowley, B.E. & Godfrey, L.R. Why all those spines? Anachronistic defences in the Didiereoideae against now extinct lemurs. *S. Afr. J. Sci.* **109**, 70–76 (2013).

22. Boisserie, J.-R., Likius, A., Vignaud, P. & Brunet, M. A new Late Miocene hippopotamid from Toros-Menalla, Chad. *J. Vertebr. Paleontol.* **25**, 665–673 (2005).
23. Lebatard, A.E. *et al.* Cosmogenic nuclide dating of Sahelanthropus tchadensis and Australopithecus bahrelghazali: Mio-Pliocene hominids from Chad. *Proc. Natl. Acad. Sci. U.S.A.* **105**, 3226–31 (2008).
24. Boisserie, J.-R., Brunet, M., Andossa, L. & Vignaud, P. Hippopotamids from Djurab Pliocene faunas, Chad, central Africa. *J. Afr. Earth Sci.* **36**, 15–27 (2003).
25. Weston, E.M. Fossil Hippopotamidae from Lothagam. in *Lothagam* (eds. Leakey, M.G. & Harris, J.M.) 441–484 (Columbia University Press, 2003).
26. McDougall, I.A.N. & Feibel, C.S. Numerical age control for the Miocene-Pliocene succession at Lothagam, a hominoid-bearing sequence in the northern Kenya Rift. *J. Geol. Soc.* **156**, 731–745 (1999).
27. Boisserie, J.R. Hippopotamidae (Cetartiodactyla, Hippopotamoidea) from Kanapoi, Kenya, and the taxonomic status of the late early Pliocene hippopotamids from the Turkana Basin. *J. Hum. Evol.* **140**, 102377 (2020).
28. Reed, K.E. Paleoecological patterns at the Hadar hominin site, Afar Regional State, Ethiopia. *J. Hum. Evol.* **54**, 743–768 (2008).
29. Campisano, C.J. & Feibel, C.S. Depositional environments and stratigraphic summary of the Pliocene Hadar Formation at Hadar, Afar Depression, Ethiopia. *Geol. Soc. Am. Spec. Pap.* **446**, 179–201 (2008).
30. Boisserie, J.-R. & White, T.D. A new species of Pliocene Hippopotamidae from the Middle Awash, Ethiopia. *J. Vertebr. Paleontol.* **24**, 464–473 (2004).
31. Harris, J.M. Family Hippopotamidae. in *Koobifora Research Project* (ed. Harris, J.M.) 31–85 (Oxford University Press, Oxford, 1990).
32. Bobe, R., Coelho, J.d.O., Carvalho, S. & Leakey, M. Early Hominins and Paleoecology of the Koobi Fora Formation, Lake Turkana Basin, Kenya. in *African Paleoecology and Human Evolution* (eds. Reynolds, S.C. & Bobe, R.) 311–331 (Cambridge University Press, Cambridge, 2022).
33. Rowan, J. *et al.* Early Pleistocene large mammals from Maka'amitalu, Hadar, lower Awash Valley, Ethiopia. *Peerj* **10**, e13210 (2022).
34. Barry, J.C. *et al.* Faunal and environmental change in the late Miocene Siwaliks of northern Pakistan. *Paleobiology* **28**, 1–71 (2002).
35. Turvey, S.T. *et al.* Late Quaternary megafaunal extinctions in India: How much do we know? *Quat. Sci. Rev.* **252**, 106740 (2021).
